# In silico proof of principle of machine learning-based antibody design at unconstrained scale

**DOI:** 10.1101/2021.07.08.451480

**Authors:** Rahmad Akbar, Philippe A. Robert, Cédric R. Weber, Michael Widrich, Robert Frank, Milena Pavlović, Lonneke Scheffer, Maria Chernigovskaya, Igor Snapkov, Andrei Slabodkin, Brij Bhushan Mehta, Enkelejda Miho, Fridtjof Lund-Johansen, Jan Terje Andersen, Sepp Hochreiter, Ingrid Hobæk Haff, Günter Klambauer, Geir Kjetil Sandve, Victor Greiff

## Abstract

Generative machine learning (ML) has been postulated to be a major driver in the computational design of antigen-specific monoclonal antibodies (mAb). However, efforts to confirm this hypothesis have been hindered by the infeasibility of testing arbitrarily large numbers of antibody sequences for their most critical design parameters: paratope, epitope, affinity, and developability. To address this challenge, we leveraged a lattice-based antibody-antigen binding simulation framework, which incorporates a wide range of physiological antibody binding parameters. The simulation framework enables both the computation of antibody-antigen 3D-structures as well as functions as an oracle for unrestricted prospective evaluation of the antigen specificity of ML-generated antibody sequences. We found that a deep generative model, trained exclusively on antibody sequence (1D) data can be used to design native-like conformational (3D) epitope-specific antibodies, matching or exceeding the training dataset in affinity and developability variety. Furthermore, we show that transfer learning enables the generation of high-affinity antibody sequences from low-N training data. Finally, we validated that the antibody design insight gained from simulated antibody-antigen binding data is applicable to experimental real-world data. Our work establishes a priori feasibility and the theoretical foundation of high-throughput ML-based mAb design.

**Highlights:**

- A large-scale dataset of 70M [3 orders of magnitude larger than the current state of the art] synthetic antibody-antigen complexes, that reflect biological complexity, allows the prospective evaluation of antibody generative deep learning
- Combination of generative learning, synthetic antibody-antigen binding data, and prospective evaluation shows that deep learning driven antibody design and discovery at an unconstrained level is feasible
- Transfer learning (low-N learning) coupled to generative learning shows that antibody-binding rules may be transferred across unrelated antibody-antigen complexes
- Experimental validation of antibody-design conclusions drawn from deep learning on synthetic antibody-antigen binding data

**Graphical abstract:** We leverage large synthetic ground-truth data to demonstrate the (A,B) unconstrained deep generative learning-based generation of native-like antibody sequences, (C) the prospective evaluation of conformational (3D) affinity, paratope-epitope pairs, and developability. (D) Finally, we show increased generation quality of low-N-based machine learning models via transfer learning.

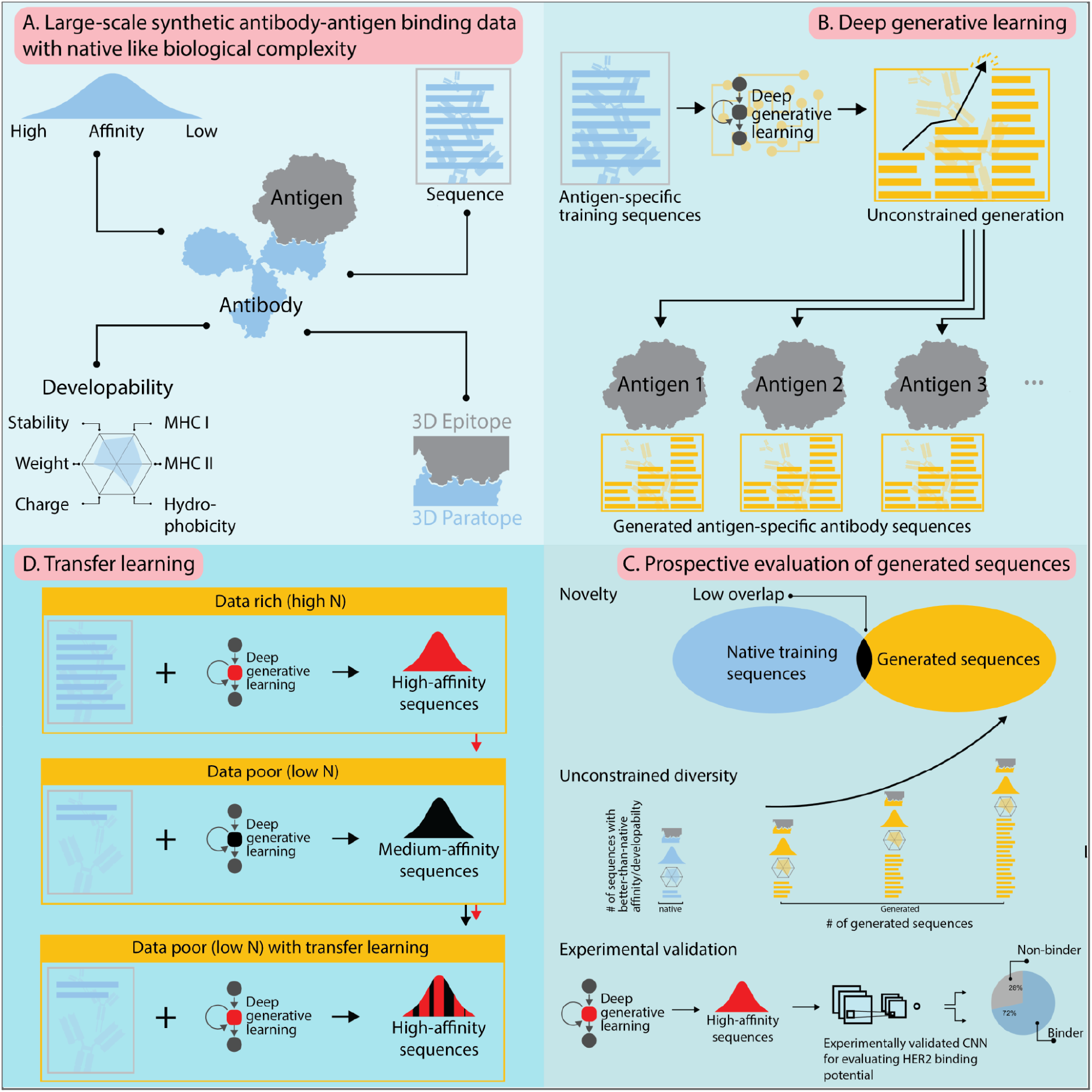

## Introduction

Monoclonal antibodies (mAbs) have proven incredibly successful in the fight against cancer and autoimmune disease (and recently, viral infections) with an estimated market size of 300 billion USD by 2025 (*1*). And recently, immense efforts to utilize mAbs for the neutralization of viral agents, such as HIV, influenza, and SARS-CoV-2 (*2–4*) are ongoing as well. So far, however, lead times to mAb discovery and design are on average >3 years (*5–8*). The reason for this is that current mAb development pipelines mostly rely on a combination of large screening libraries and experimental heuristics with very little to no emphasis on rule-driven discovery (*9*). Recently, it has been increasingly postulated that machine learning (ML) may be useful in accelerating antibody discovery – especially when applied to large-scale antigen-specific display library screening antibody sequencing data (*10*). However, formal proof that ML can generate antibody sequences that are 3D-antigen-specific (affinity, paratope, epitope) if only provided with 1D-sequence training data (the most abundant class of available antigen-specific antibody data) alone is missing.

Recent reports suggest that ML may be able to learn the rules of efficient antibody (protein) design (*6, 10–17*). Specifically, Amimeur and colleagues (*18*) trained generative adversarial networks (GANs) (*19*) on sequences obtained from the Observed Antibody Space (OAS) database (*20*) to demonstrate the capacity of deep generative networks to discover mAbs with certain developability parameters. Friedensohn and colleagues (*21*) trained variational autoencoders (VAE) (*22*) on BCR-sequence data to identify antibody sequences in cohorts of mice immunized with various antigens and to generate novel antigen-binding sequences. Widrich et al. as well as Davidsen et al. have used LSTMs/VAEs to generate TCRβ sequences with the aim to generate realistic immune repertoires (*23, 24*). Finally, Eguchi et al. built class-specific backbones using VAE to generate 3D coordinates of mAbs (*25*). However, while several generative deep learning methods have been explored for the in silico generation of immune receptor sequences, these strategies did not allow for the examination of whether the generated sequences follow the same antigen-specificity distribution as the input training data. This is due to the absence of large-scale antigen-annotated antibody sequence training data and the lack of high-throughput techniques for validating antigen binding of ML-generated antibody sequences.

Here, we investigated whether generative deep learning can learn 3D-affinity and epitope information from 1D antibody sequence data. This was done by using two oracles (external validators): (i) an *in silico* framework that enables unrestricted validation (*prospective evaluation*) of the biological activity (paratope, epitope, affinity) of generated antibody sequences. Specifically, we used an *in silico* antibody-antigen binding simulation framework (which respects the biological complexity of antibody-antigen binding to the largest extent possible), called *Absolut!* (*26*) that can annotate large collections of antibody sequences with synthetic binding affinities (specificity) to a synthetic 3D-antigen, which allows the assembly of large-scale complete-knowledge training data (*27, 28*). Due to its ability to annotate newly generated antibody sequences with antigen-binding information, Absolut! resolves the current problems of *in silico* validation of generated sequences (*29, 30*). And (ii) an experimentally validated deep learning classifier that was trained on binders and non-binders to epidermal growth factor 2 (HER2) (*31*). Our work provides the basis for the ML-driven design of fit-for-purpose antibodies with respect to binding affinity, epitope, and developability.

## Results

### Deep learning generates novel antigen-specific CDR-H3 sequences across a wide range of developability parameters

ML-based generation of new antibody sequences with desired biological properties requires large experimental datasets and a method to test the generated sequences for such properties. To address the absence of large experimental antigen-specific antibody-antigen datasets to train and test deep antibody generative models, we leveraged Absolut!—a software suite that simulates the binding of antibody sequences to 3D antigens. Absolut!, to a large extent, replicates and recaptures the biological properties and complexity of experimental antibody-antigen binding (*26*). We utilized our previously published dataset of seven million (7×10^6^) native antibody (CDR-H3) amino acid sequences (*33*) (see Methods) and (via Absolut!) computed their binding to 10 protein antigens (Table 1, Figure 1A). Briefly, synthetic antibody-antigen complexes were obtained by (i) iterating over all possible binding positions between a sequence and an antigen to find the optimal binding position and (ii) by calculating the resulting binding affinity, paratope, epitope, and structural fold for each antibody CDR-H3 sequence (see *Methods*). Following affinity annotation, a set of six developability parameters (Table 2) was calculated for each CDR-H3 sequence (Figure 1A). CDR-H3 amino acid sequences equipped with affinity, paratope, epitope, and developability information are henceforth termed *antigen-annotated* CDR-H3 sequences.

**Figure 1.**
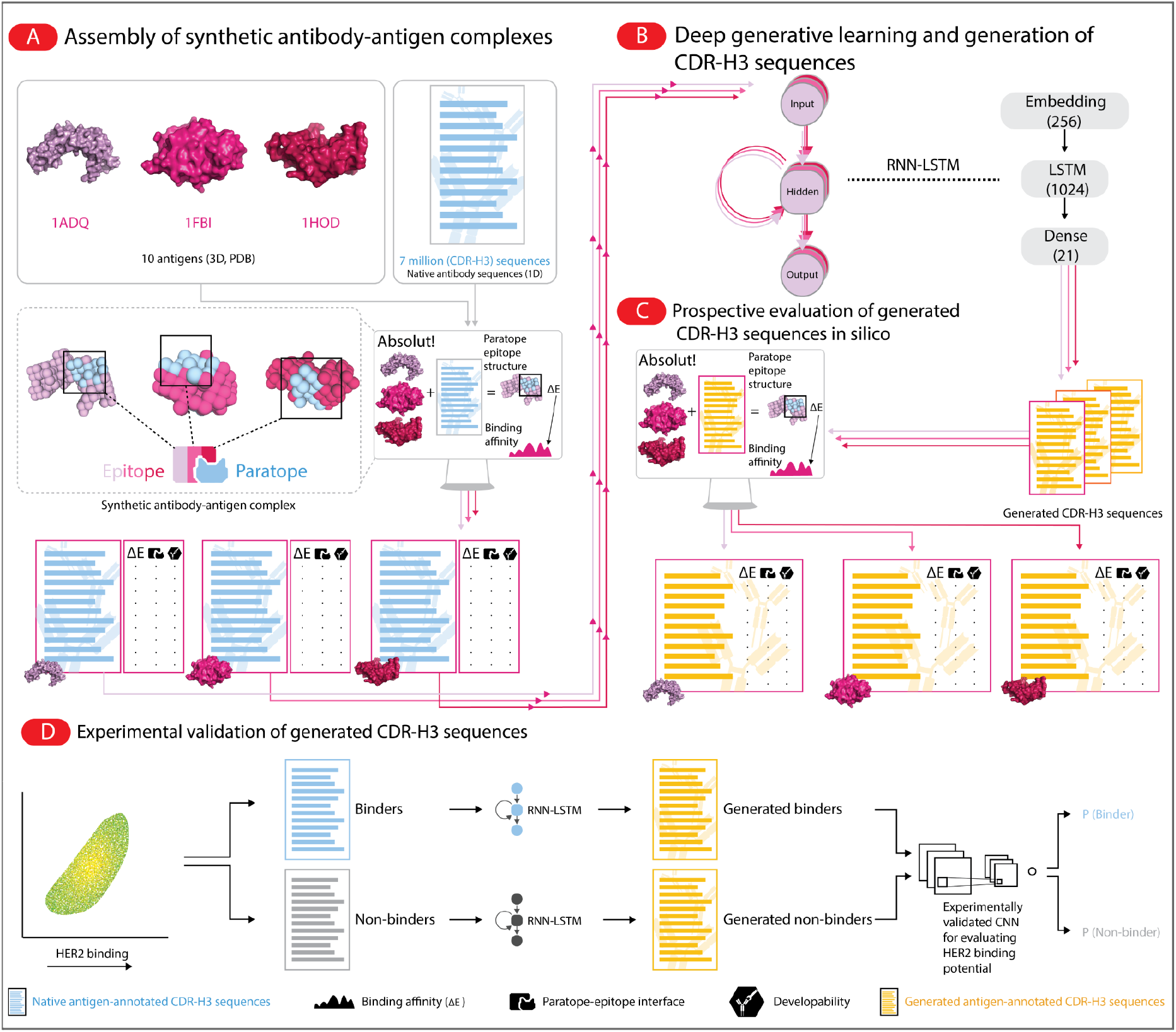
Workflow for ML-based antibody design and evaluation thereof. **(A)** Generation of in silico training datasets with binding paratope, epitope, and affinity annotation. Briefly, PDB 3D antigen structures were obtained from the Antibody Database (*32*) and native antibody sequences (CDR-H3) were obtained from Greiff and colleagues (*33*). CDR-H3 sequences were annotated with their corresponding affinity and epitope to each antigen using the Absolut! software suite (*26*). In addition, six widely used developability parameters were calculated for each CDR-H3 sequence (see Table 2). **(B)** Training a generative model on high-affinity CDR-H3 sequences to each antigen. Native linear 1 D antigen-specific CDR-H3 sequences were used as input to train sequence-based RNN-LSTM generative models. Of note, the RNN-LSTM architecture did not receive any explicit 3D-information on the paratope, epitope, affinity, nor the developability of a given sequence. **(C)** Large-scale in silico CDR-H3 sequence generation and binding validation. Following training, the deep models were used to generate new CDR-H3 sequences, which were then evaluated (prospectively tested) for their antigen-specificity (affinity, paratope, epitope) using Absolut! (simulation) and annotated with developability-associated parameters. **(D)** To validate our RNN-LSTM model beyond the binding simulation setting, we trained the model with experimental HER2 binder and non-binder CDR-H3 sequences and generated 6×10^6^ new sequences, and evaluated the generated sequences against an experimentally validated CNN-based HER-2 classifier model as reported in (*31*).

**Table 1.** List of 3D-antigens used in the deep-learning-based antibody generation pipeline.

| PDB-ID | Antigen | Species of origin |
| --- | --- | --- |
| 1ADQ | IGG4 REA FC<br>(206 residues, ~ 22kDa) | Homo sapiens |
| 1FBI | GUINEA FOWL LYSOZYME<br>(129 residues, ~14kDa) | Numida meleagris |
| 1H0D | ANGIOGENIN<br>(122 residues, ~ 13kDa) | Homo sapiens |
| 1NSN | STAPHYLOCOCCAL NUCLEASE<br>(138 residues, 15kDa) | Staphylococcus aureus |
| 1OB1 | MAJOR MEROZOITE SURFACE PROTEINS MSP1-19<br>(95 residues, ~10kDa) | Plasmodium falciparum |
| 1WEJ | CYTOCHROME C<br>(104 residues, ~11kDa) | Equus caballus |
| 2YPV | MENINGOCOCCAL VACCINE ANTIGEN FACTOR H<br>TITLE 2 BINDING PROTEIN (229 residues, ~25kDa) | Neisseria Meningitidis |
| 3RAJ | ADP-RIBOSYL CYCLASE 1<br>(230 residues, ~25kDa) | Homo sapiens |
| 3VRL | GAG PROTEIN<br>(73 residues, ~8kDa) | Human Immunodeficiency<br>Virus 1 |
| 5E94 | GLUCAGON-LIKE PEPTIDE 1 RECEPTOR<br>(110 residues, ~ 12kDa) | Homo sapiens |

\*Weight (in kilodaltons, kDa) was estimated by using the average amino acid weight of 110 Da. Missing residues were omitted.

**Table 2.** Antibody developability parameters and their computational implementation.

| Developability parameter | Computational descriptor | Computational tool (function) |
| --- | --- | --- |
| Charge | charge at pH 7 | `Bio.SeqUtils.ProtParam`<br>(`charge_at_pH`) (59) |
| Hydrophobicity | Gravy | `Bio.SeqUtils.ProtParam`<br>(`gravy`) (59) |
| Stability | Instability index | `Bio.SeqUtils.ProtParam`<br>(`instability_index`)<br>(59) |
| Affinity to MHC class II molecules | Average rank of predicted affinity to MHC II molecules | NetMHCIIpan 4<br>(60) |
| Affinity to MHC I molecules | Average rank of predicted affinity to MHC I molecules | NetMHCpan 4 (60) |
| Weight | Molecular weight (kDa) | `Bio.SeqUtils.ProtParam`<br>(`molecular_weight`) (59) |

We examined the capacity of a deep (autoregressive) generative model (recurrent neural networks with long short-term memory—RNN-LSTM) to generate (design) novel antigen-specific sequences as follows. (i) We trained the RNN-LSTM model on antigen-specific CDR-H3 sequences (top 1% affinity sorted sequences, n_seq_=70 000) (Figure 1). Importantly, we did not provide explicitly the affinity or paratope/epitope information in the training process. (ii) Subsequently, we used the trained model to generate new CDR-H3 sequences (n_seq_=70 000) (Figure 1), (iii) which we evaluated in terms of antigen specificity (using Absolut!), sequence novelty, and developability (Figure 1C).

The binding affinity (Figure 2A), as well as the paratope fold and epitopes of generated CDR-H3 sequences, mirrored very closely those of the native (training) CDR-H3 sequences (Figure 2B). Novel paratope folds and epitopes were also discovered as observed by the paratope fold and epitope diversity (*34*) of generated CDR-H3 sequences that were higher than those of native (training set) sequences (Figure 2B). The within-sequence similarity, as measured by the distribution of Levenshtein distance (LD) between CDR-H3 sequences within the set of native or generated CDR-H3 sequence datasets was preserved (Figure 2D, Fig. S4) as were long-range sequence dependencies (gapped k-mer decomposition, Pearson correlation 0.864–0.907, Figure 2C). To exclude the possibility that generated CDR-H3 sequences showed high affinity merely by virtue of their similarity to the training input, we validated that the generated CDR-H3 sequences were novel (<1% overlap between generated and native antigen-specific sequences, Figure 2E) both measured by exact sequence identity or based on sequence similarity (median Levenshtein distance between generated and native CDR-H3 sequences: ≈9–10 a.a., Figure 2D). Thus, deep generative learning explores non-trivial novel sequence spaces. We excluded the possibility that the chosen RNN-LSTM-architecture would be biased to generate high-affinity sequences as default by training on CDR-H3 sequence sets that contain (*1*) exclusively low-affinity CDR-H3 sequences [generating exclusively low-affinity CDR-H3 sequences] and (ii) CDR-H3 sequences spanning the entire affinity spectrum (generating CDR-H3 spanning the entire affinity spectrum) (Fig. S3). Finally, the distribution of developability parameters of generated CDR-H3 sequences largely mirrored but also expanded the range of parameters of native antigen-specific sequences (Figure 2F).

**Figure 2.**
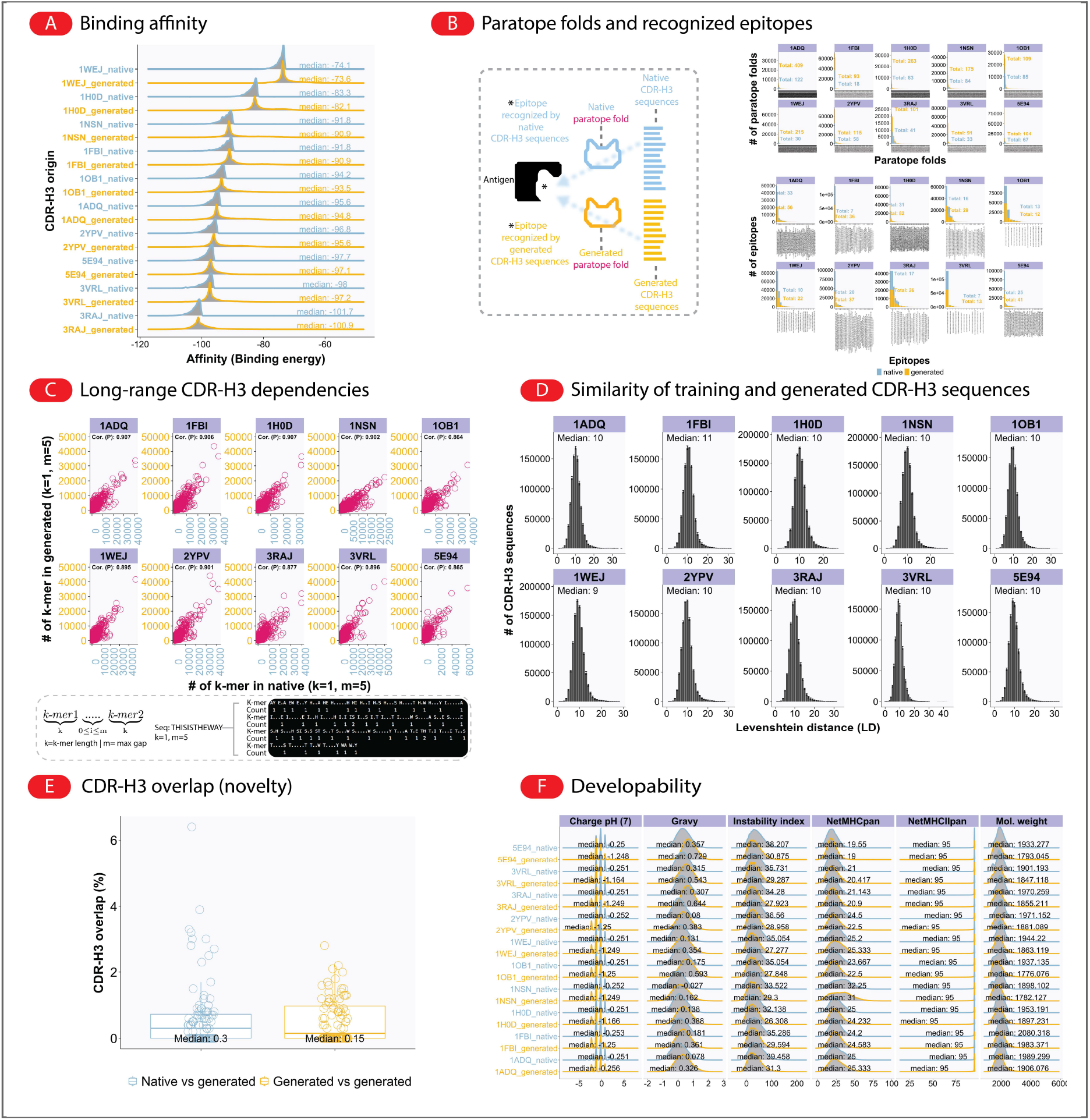
Deep learning (RNN-LSTM) generates novel antigen-specific CDR-H3 sequences across a wide range of developability parameters. **(A)** Comparison of training and generated affinities. The affinity of training antigen-specific CDR-H3 sequences (n_seq_=70 000, blue) to 10 different 3D antigens obtained from PDB (see Table 1). The affinity of the 70 000 generated CDR-H3 sequences from the 10 RNN-LSTM models was shown in yellow. **(B)** Comparison of training and generated sequences for epitope recognition. Absolut! was used to compute the (A) affinity and (B) paratope fold/epitope of the training data (see Methods: Generation of lattice-based antibody-antigen binding structures using Absolut!). **(C)** Pearson correlation (range: 0.864–0.907) of CDR-H3 sequence composition between training (“native”) and generated datasets quantifying the preservation of long-range dependencies. CDR-H3 sequence composition was measured using gapped k-mers where the size of the k-mer was 1 and the size of the maximum gap varied between 1 and 5. **(D)** CDR-H3 sequence similarity (Levenshtein distance, LD) distribution determined *among* training and generated CDR-H3 sequence datasets (see Fig. S4 for the LD distribution of CDR-H3 sequences with the native and generated set, respectively). **(E)** CDR-H3 sequence novelty (overlap) defined as CDR-H3_antigen_x_⋂CDR-H3_antigen_y_/70 000, where x and y are the 10 antigens listed in Table 1) of CDR-H3 sequences (median overlap <0.5% → novelty: >99.5%) between both “native and generated” and “generated and generated” datasets across all antigen combinations. **(F)** Developability parameter distribution between training and generated CDR-H3 sequences overlaps substantially (see Table 2 for a description of developability parameters used).

### On-demand generation of large amounts of CDR-H3 sequences with broad developability and affinity that match or exceed the training sequences

Following the observation that deep generative models were capable of generating novel CDR-H3 sequences that mirror very closely the binding and developability properties of native CDR-H3 sequences (Figure 2), we hypothesized that such models are useful for generating large quantities of CDR-H3 sequences with similar or better affinities than those of the native ones. To assess this hypothesis, (i) we grouped the native antigen-specific CDR-H3 sequences (n_seq,training_= 70 000, top 1%) into four affinity categories based on their binding energy (low energy → high affinity): ultimate binder (max native–⅓), penultimate binder (⅓–⅔), binder (⅔–min native), and hyperbinder (affinity>native max, i.e., higher affinity than found in the training data CDR-H3 sequences, see schematic in Figure 3A), and (ii) we generated, for each antigen, 7×10^5^ unique antigen-specific CDR-H3 sequences (i.e., 10 times larger than the training dataset), and evaluated the generated CDR-H3 sequences with respect to the four categories. Broadly, we found that the number of binders in all four categories increased as the generated sequences increased (Figure 3A). Specifically, when the number of generated CDR-H3 sequences equaled that of the training data (n_seq,generated_=70 000), we found binders in the same order of magnitude in all categories of binders (binder–ultimate binder) compared to the native (training) dataset (blue lines), except for hyperbinders (as the native populations have, per definitionem, no hyperbinders). At n_seq,generated_=7×10^5^, the quantities of discovered binders far eclipsed those of the native binders in all four categories by ~4-fold (Figure 3A) suggesting that generative learning may be used for a highly exhaustive discovery of novel binders. Importantly, the discovery of CDRH3 sequences with superior predicted binding affinity compared to the native sequences (hyperbinder) further illustrates the importance of deep generative models in the design and discovery of high-affinity CDR-H3 sequences (*11, 18, 21, 35*). Hyperbinders showed affinity improvements over native CDR-H3 sequences in the range of 0.4–4.4% (percentages were calculated against the maximum affinity [lowest energy] of each antigen’s training dataset) and median LD (against native binders) of 10 to 14. To summarize, our RNN-LSTM models were able to generate large quantities of non-redundant CDR-H3 sequences that match or exceed the affinity of the training sequences.

**Figure 3.**
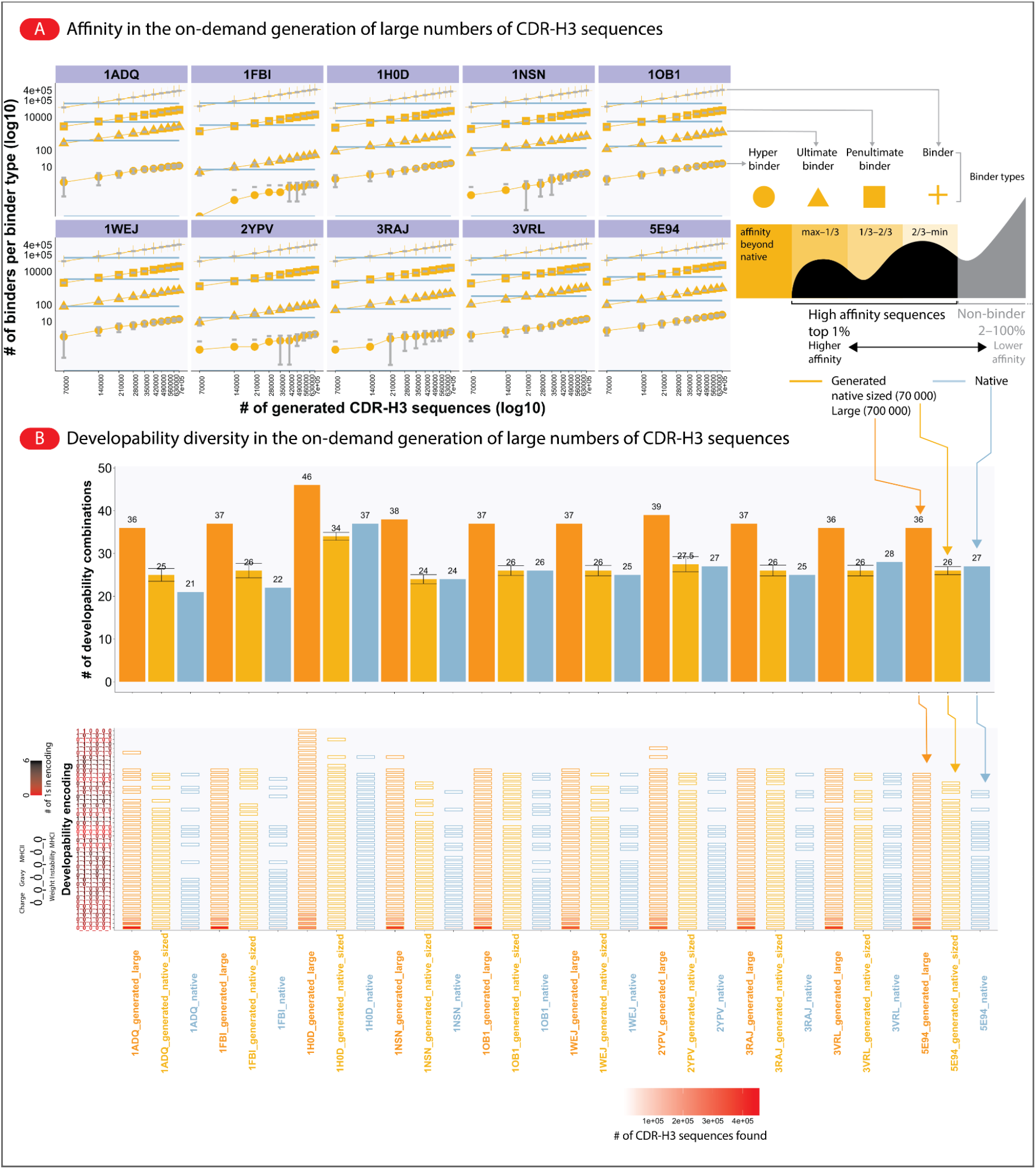
Exhaustive generation reveals better antibodies than are present in the training dataset. **(A)** To examine the ability of the RNN-LSTM model to generate CDR-H3 sequences beyond the native realms (in terms of quantity and affinity), we first binned the native high-affinity antigen-specific training CDR-H3 sequences into four affinity classes: hyperbinder (affinity >max native), ultimate binder (max native>–1/3), penultimate binder (1/3–2/3), and binder (2/3–min native). Following binning, we used deep generative models to generate 700K new sequences, devised 10 cutoffs in the increment of 70K (70K[native sized], 140K…700K[large]), subsampled 10 times (from the 700K generated sequences) and counted the number of novel sequences in each cutoff. Native and generated sequences are shown in blue and yellow; error bars are shown for subsampled sequences. We found that, for all affinity classes, the number of sequences in each class increases as a function of the total number of generated sequences. In addition, we found sequences that possess a higher affinity than the native-training sequences (called hyperbinders) with affinity improvements over native CDR-H3 sequences ranging between 0.04–4.4% [depending on the antigen, percentages were calculated relative to the minimum affinity per antigen]. **(B)** To examine the diversity and preferences of developability combinations, we annotated each CDR-H3 sequence with a binary developability encoding. Briefly, we binned each developability parameter in two bins (low=min–median and high=median–max) and annotated each sequence with a composite binary encoding from all six developability parameters (i.e., 0_0_0_0_0_1 indicates that the sequence has a low charge, low molecular weight, low gravy index, low instability index, low affinity to MHCII and high affinity to MHC). We found that the generated CDR-H3 sequences yielded larger ranges of developability combinations in *native-sized* generation (n_seq_=70 000) and *large* generation (n_seq_= 7×10^5^). Error bars indicate the standard deviation for the subsampling.

In the same vein, we hypothesized that deep generative models would prove useful for generating CDR-H3 sequences with similar or richer developability profiles to native CDR-H3 sequences (higher number of combinations or constraints on developability parameter values). To this end, we devised a binary developability encoding wherein each developability parameter (Table 2) is grouped into two categories: *low* (parameter values that range between the min and median of the distribution of the parameter) and *high* (parameter values that range between the median and max of the distribution of the parameter) and annotated each CDR-H3 sequence with a composite developability encoding combining all six developability parameters here examined (Figure 3B). For instance, the encoding 0_0_0_0_0_l indicates that the thus annotated CDR-H3 sequence has a *low* charge (*0*), *low* molecular weight (0), *low* gravy index (0), *low* instability index (0), *low* affinity to MHCII (0) but a *high* affinity for MHCI (1). Subsequently, we compared the total number of developability parameter combinations populated by the generated sequences (against native sequences) in two conditions: *native-sized* wherein the number of generated sequences matches the number of sequences in the native (training) dataset (n_seq,generated_=70 000) and *large* where the number of generated sequences is an order of magnitude larger than the native training sequences (n_seq,generated_=7×10^5^). We observed a larger number of developability parameter combinations in the generated populations (Figure 3B). Specifically, *native-sized* generation yielded 29–39 developability parameter combinations (45–61% of all possible combinations), *large* generation yielded 33–44 (52-69% of all possible combinations) as compared to native sequences that yielded 21–37 combinations (33–58% of all possible combinations). Pearson correlation between the counts of developability parameter combinations in native and generated sequences was high (Pearson cor: 0.74–0.99, Figure 3B). In other words, deep generative models can be leveraged to generate antibody sequences that are equally or more diverse than native (training) ones in terms of developability profile.

### The quality of ML-based antibody sequence generation is function of the size of the training data

The absence of large antigen-annotated antibody sequences and structural datasets remains a major challenge in developing robust machine learning methods for antibody-antigen binding prediction as well as antigen-specific generation of monoclonal antibodies (*11, 36*). Furthermore, the precise amount of antibody sequence data necessary to recover native-like antibody affinity, epitope, and developability is a subject of ongoing investigations (*10–12*). Therefore, within the framework of our simulation suite Absolut!, we examined how the number of training CDR-H3 sequences impacts the resulting binding affinity of the generated CDR-H3 sequences. To this end, from the top 1% antigen-specific CDR-H3 sequences (n_seq_=70 000), we created smaller datasets of antigen-specific CDR-H3 sequences (n_seq,subsample_=700, 7000, 10 000, 20 000, 30 000, 40 000, 50 000, 60 000, and n_replicates_=5); trained deep generative models on these subsets; and compared the resulting binding affinity against native CDR-H3 sequences and CDR-H3 sequences generated by models trained on the top 1% (nseq=70 000) antigen-specific CDR-H3 sequences (called “data-rich” model). We found that the correspondence between the binding affinity and epitope recognition of native and generated CDR-H3 sequences increased as a function of the number of training CDR-H3 sequences (Figure 4A, Fig. S5). Specifically, our models recovered very closely the native affinity (as measured by median energy) when we used 20 000 or more training CDR-H3 sequences (Fig. S5). Similarly, the agreement of epitope occupancy between generated CDR-H3 sequences increases as a function of the sequence size of the training set (Fig. S8, Fig. S11). Of note, we found that the agreement of epitope occupancy between native and generated CDR-H3 sequence was already reasonable at a small training dataset (ntrain=700) for antigens with fewer epitopes (e.g., 3VRL, see Figure 2B bottom panel) (Fig. S8, Fig. S11). In contrast, antigens with more epitopes (e.g., 1H0D, Figure 2B, bottom panel) required larger training datasets for reaching a high concordance with the epitope occupancy observed in the training dataset (Fig. S8, Fig. S11).

**Figure 4:**
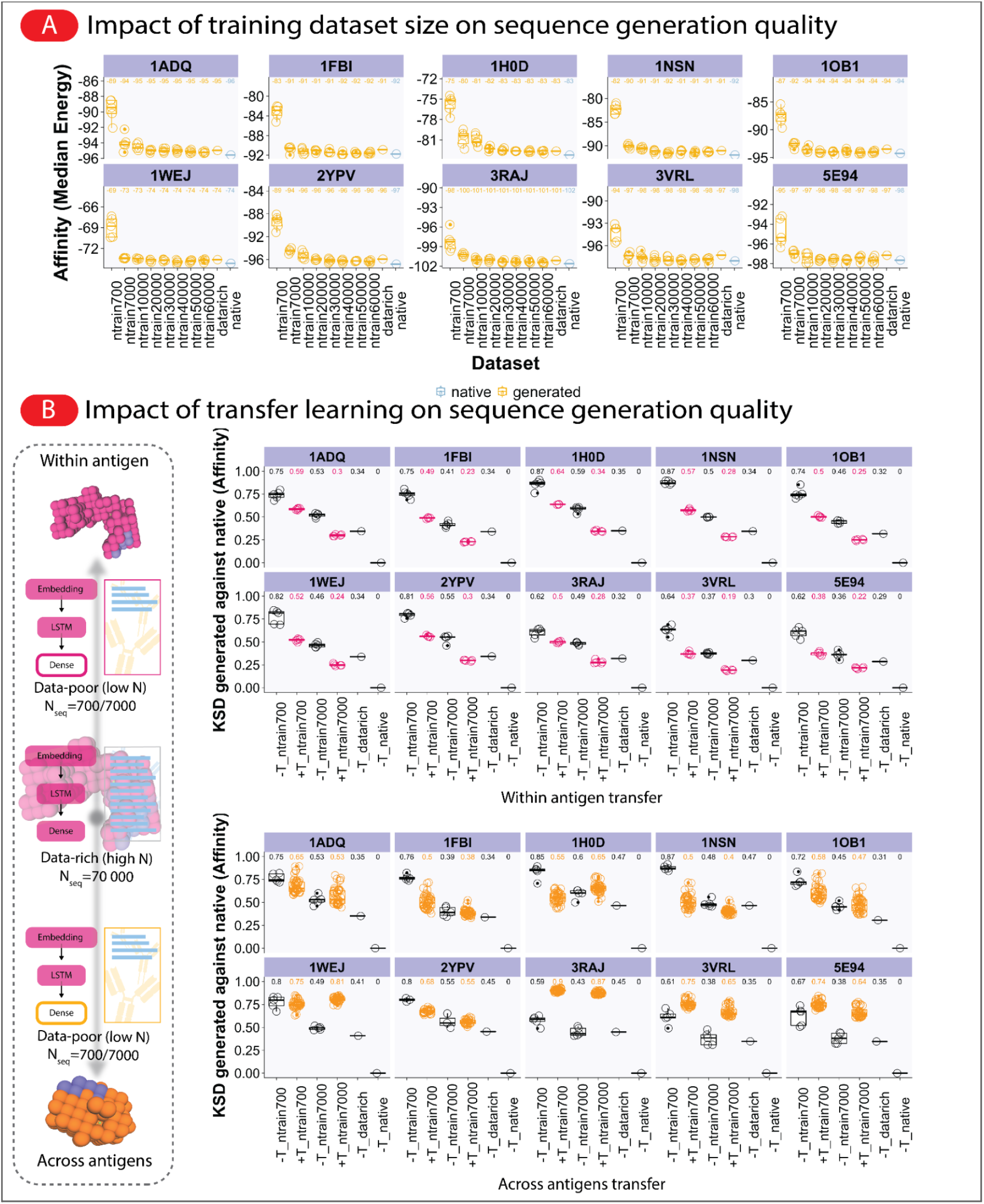
Generation quality of antibody sequences depends on the size of the training dataset and transfer learning enables the generation of higher-affinity CDR-H3 sequences from lower-sized training datasets. **(A)** To examine the impact of sample size on the resulting binding affinity and epitope (see Fig. S8) of generated CDR-H3 sequences, we created smaller training datasets (n_seq,subsample_=700, 7000, 10 000, 20 000, 30 000, 40 000, 50 000, 60 000, and n_replicates_=5) from the full antigen-specific CDR-H3 sequences (n_seq, training_=70 000), trained deep generative models on the subsets and compared the binding affinity and epitope against affinity and epitope from models trained on the full data and the native affinity and epitope (see Fig. S8 for correlations of CDR-H3 epitope occupancy). We found that models trained on the larger dataset sizes (>2×10^4^), but not the smaller subsets (in the order of 10^3^ or 10^2^), sufficiently replicate the distribution of binding affinity and epitope CDR-H3 sequences. **(B)** To investigate whether transfer learning may be used to improve the affinity and epitope (see also Fig. S9–Fig. S13) binding of CDR-H3 sequences generated by models trained on smaller-sized datasets, we constructed a transfer architecture wherein embedding and RNN-LSTM layers from a “data-rich” model (high N, n_seq, training_=70 000) were stacked atop of a fresh dense layer and training the resulting ‘transfer’ model on lower-sized datasets (data-poor; low N, n_seq,training_=700/7000). Two types of transfer experiments were performed: a within-antigen transfer experiment (e.g., between a data-rich model of an antigen *V* and data-poor models of the same antigen *V*) and a between-antigens (across antigens) transfer experiment (e.g., between data-rich model of an antigen *V* and data-poor model of antigen *G*). We used Kolmogorov–Smirnov distance (KSD, range: 0 for identical distribution, increasing value for increasing dissimilarity between distributions) to quantify the similarity between affinity distributions of CDR-H3 sequences generated by the models with transfer learning (+T) and without transfer learning (-T). Smaller KSD values indicate that the compared affinity distributions are similar and a larger value signifies dissimilarity of affinity distributions. For within transfer experiments, we found marked reductions of KSD values (against the native population) in all antigens signifying the transferability of general antibody-antigen binding features within antigens. For across-antigens transfer experiments, 7 out of 10 antigens showed reductions in KSD values in at least one transfer scenario (n_seq, training_=700 or 7000, Figure 4B) suggesting the transferability of antibody-antigen binding features across antigens.

In summary, 20 000 CDR-H3 sequences were sufficient to train models that reproduce native-like affinity. We note that our simulation framework Absolut! does not operate at atomistic resolution (*26*), thus, n_seq,training_ in the order of 20 000 shall only be regarded as a lower bound of the number of training CDR-H3 sequences necessary to train a robust deep generative model, in comparison with a higher dimensionality of binding modes in experimental datasets.

### Transfer learning enables the generation of high-affinity CDR-H3 sequences from lower-sized (low-N) training datasets

Based upon the observation that lower-sized training datasets failed to produce CDR-H3 sequences with native-like binding affinity and epitope binding, we asked whether the generation quality of models trained on lower-sized datasets (data-poor, “low-N, (*37*)”, n_seq,training_=700 and 7000) may be improved by transferring learned features from models trained on larger training datasets, which were found to be sufficient for achieving a native-like affinity (data-rich, n_seq,training_=70 000, Figure 2, Figure 4A). We examined this question by constructing a transfer learning architecture wherein pre-trained embedding and RNN-LSTM layers from a data-rich model were stacked atop of a new fully-connected layer with the resulting “transfer” model subsequently being trained on lower-sized datasets (Figure 4). We performed two different transfer learning experiments termed (i) *within antigen* and (ii) *across antigens* transfer. (i) *within antigen* transfer describes a transfer experiment involving the same antigen (this transfer setting serves as control for the functioning of the transfer architecture). That is, pre-trained embedding and LSTM layers from a data-rich model based on CDR-H3 sequences specific for an antigen *V* were stacked atop of a new dense layer; the resulting architecture was trained on lower-sized datasets (n_seq,training_=700 and 7000) of antigen *V* In contrast, (ii) *across antigen* transfer identifies a transfer experiment involving different antigens, e.g., a data-rich model of an antigen *V* and data-poor (lower-sized datasets n_seq,training_=700 and 7000) models of antigen *G* (see Methods and Figure 4B). Following training, for each antigen, we generated a total of 100 000 CDR-H3 sequences (10 000 sequences, 10 replicates) and measured the generation quality with respect to affinity and epitope. We used the Kolgomorov–Smirnov distance (KSD) to quantify the similarity between the generated binding affinity distributions and the native affinity distribution. A small KSD indicates that the compared affinity distribution is similar and increasing KSD indicates increased dissimilarity. We observed marked reductions of KSD values (against the affinity distribution of the native population) for the within antigen transfer in all models (Figure 4B, upper panel), which signifies the availability, learnability, and transferability of general antibody-antigen binding features within an antigen. For the across antigens transfer experiments, 7 out of 10 antigens showed reductions in KSD values in at least one transfer scenario (n_seq,training_=700 or 7000, Figure 4B, lower panel) suggesting the transferability of antibody-antigen binding features across antigens and the multi-faceted nature of the signal per antigen learned (nota bene, the medians of binding affinities in the across antigens transfer scenario were closer to the native and data-rich affinities in all 10 antigens, Fig. S6). For epitope similarity, we used Pearson correlation (Fig. S9–Fig. S10) and overlap (Fig. S12–Fig. S13) to quantify the concordance between epitopes recognized by native and generated CDR-H3 sequences. Similar to affinity, we found better concordance both for the within and across antigens transfer (increasing Pearson correlation values, Fig. S9 and Fig. S10). Interestingly, the number of recognized epitopes jumped in across-antigens transfer (Fig. S13) whereas in the within-antigen transfer (Fig. S12) the number of recognized epitopes dropped, hinting at the utility of across antigens transfer in generating epitope diversity. In summary, our in silico experiments suggest that transfer learning may represent a suitable method for generating high-affinity CDR-H3 sequences from lower-sized training datasets.

### Experimental validation of antibody-design conclusions drawn from ML training on simulated antibody-antigen binding data

Experimental validation at the scale of the number of antibody sequences that can potentially be ML-generated (Figure 3) remains an unresolved technological problem. One solution to this challenge are experimentally validated ML-classifiers that can screen the potential sequence space for binders. One such classifier for HER-2 binders was previously developed by Mason and colleagues (*31*). Briefly, this CNN-based classifier classifies CDR-H3 amino acid sequences for their potential to bind HER2; all sequences annotated with a binding probability of p>0.5 are considered binders. Mason and colleagues validated this classifier experimentally by the expression and testing for binding of both predicted binders and non-binders. Given that the experimental system by Mason et al. is very similar to the one simulated in this work, i.e., testing of binding of CDR-H3 sequences, we concluded that the CNN-classifier can be used to evaluate the experimental binding potential of the output of our RNN sequence generator (Figure 1B). Therefore, we performed the following experiment: we trained separate RNN-LSTMs on the experimentally verified 11300 HER-2 binders (“RNN-LSTM binder model”) and the 27539 HER-2 non-binders (“RNN-LSTM non-binder model”) of the Mason dataset and used the models to generate 6×10^6^ sequences in each case (Figure 5A). These generated sequences were assessed for their HER2-binding potential using the experimentally verified CNN-classifier described above (Figure 5A). We found that 72% of the generated CDR-H3 sequences from the RNN-LSTM binder model were scored as binders and we ascertained that the generated sequences follow the positional amino acid dependencies (usage) of the experimentally verified training data (low mean-squared error, MSE, Figure 5B). We verified that these results were not random by CNN-scoring the CDR-H3 sequences from the RNN-LSTM non-binder model where only 14% of the generated CDR-H3 sequences were scored as binders. Thus, we validated that the RNN-LSTM trained on experimentally-determined HER-2-binding sequences, successfully generated sequences classified as HER-2-binders by the experimentally validated CNN and that a training dataset in the order of 1–2×10^4^ sequences (as also observed with our synthetic data, Figure 4A) is sufficient to generate CDR-H3 sequences that bind the target antigen. Of note, these results also suggest that de novo antibody design is feasible using only binding sequences (positive data) for ML model training.

**Figure 5.**
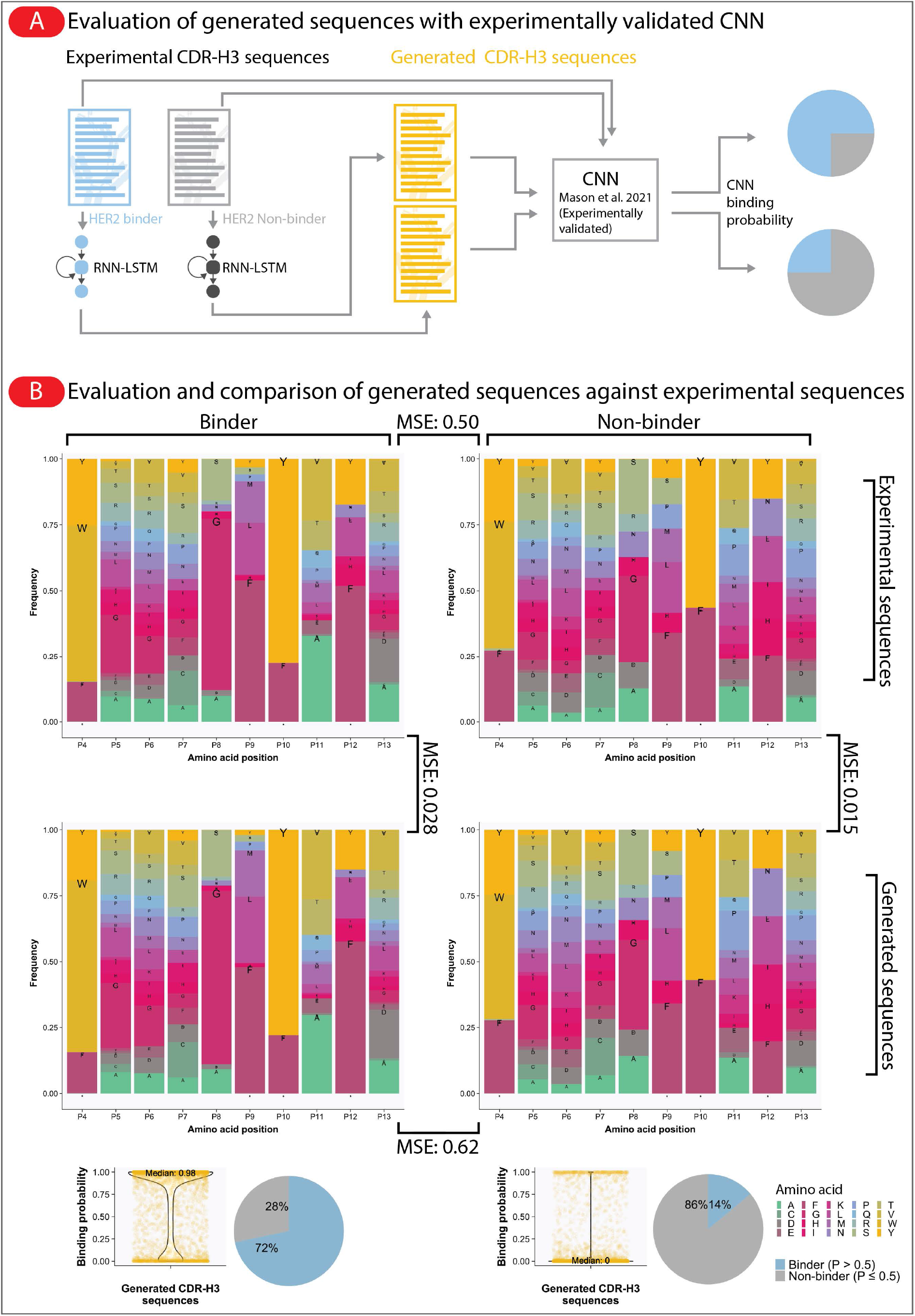
RNN-LSTM model trained on experimentally validated binders (not synthetic sequences) generated native-experimental-like binders. **(A)** To validate that our model can not only reproduce properties of native-like *synthetic* sequences of binders but also experimentally determined binders, we trained the model with binders and non-binders obtained from recently published experimental data of binders against human epidermal growth factor 2 (HER2) (*31*), generated 6×10^6^ sequences and scored the sequences with the Mason et al. CNN classifier (the CNN classifier outputs a probability value between 0–1. CDR-H3 sequences with probability >0.5 are categorized as binders). **(B)** We found that a large proportion (72%) of generated sequences originated from the model that was trained on HER2 binders had high prediction probability values (P>0.5, as used in Mason et al.) and correspondingly the majority (84%) of the generated sequences originated from the model that was trained on HER2 non-binders had low prediction probability values (P≤0.5) as well as similar amino acid usage across positions as shown by the amino acid frequency plots (as described in the Mason et al. study). Of note, the mean squared error (MSE) values of the generated sequences were comparable to the values of experimental sequences indicating that our generated sequences preserve the characteristics of the experimental sequences (HER2-binders, HER2-non-binders) rather well.

## Discussion

We have here provided the in silico proof-of-principle that deep learning can learn the rules of 3D-antibody-antigen interaction from 1D antibody sequence data alone by showing (in a 3D-lattice space) that novel antibody variants with high affinity and specific epitope binding can be generated based on sufficiently large training data (Figure 2 and 4). Among the generated antibodies, for all tested antigens (10 out of 10), we detected novel antibody sequences that exceeded in affinity those found in the training dataset (Figure 3). ML-based sequence generation also allowed for the discovery of novel developability parameter combinations (Figure 3). For the machine learning model used, we determined the number of training CDR-H3 sequences necessary (>2×10^4^) for generating high-affinity CDR-H3s and demonstrated that these numbers may be reduced by transfer learning (Figure 4). Finally, we experimentally validated the antibody-design conclusions drawn from ML training on simulated antibody-antigen binding data (Figure 5). More broadly, while the primary objective of this paper was the proof-of-principle study of antibody generative learning, the secondary objective was to develop a set of analytical approaches that may help study the quality of generated antibody sequences in future studies with a similar aim.

In this work, we chose an RNN-LSTM based language modeling approach as it represents a competitive baseline to the state-of-the-art transformer-based architecture (*38*). Recently, variational autoencoders (VAE), as well as generative adversarial networks (GAN), have also been used for generating T and B-cell receptor sequences (*18, 21, 24*). However, both in the area of natural language processing as well as in the area of generative models for small molecules, GANs and VAEs remain less competitive (Semeniuta et al., 2018; Preuer et al., 2018). Although we decided to use an RNN-LSTM as a generative model, we hypothesize that any accurate language model, for example, transformer architectures (*26*), would procure similar results and conclusions. Further benchmarking is needed in the area of generative protein design.

A common problem with deep learning-generated sequence data is that methods may reproduce the training data with minimal changes, which has been termed the “copy problem” by Renz and colleagues (*39*). The copy problem is especially prevalent when the capacity for high-throughput testing of molecular properties (in our case, antigen binding and developability) is unavailable. The absence of prospective testing capacity precludes the functional (e.g., antigen binding) evaluation of the generated dataset, which renders addressing the copy problem somewhat unfeasible (merely testing sequence diversity on sequences of which the binding mode is unknown does not elucidate the extent of diversity for a given binding mode for example). In this work, we were able to address and exclude a copy problem by evaluating all generated sequences for *both* binding as well as for sequence diversity (Figure 2–Figure 4).

The transfer learning experiments demonstrated the capacity of deep learning models trained on large collections of CDR-H3 sequences to augment weaker datasets (smaller datasets that fail to reproduce faithfully the affinity and epitope of native sequences) for both within and across antigens scenarios (Figure 4). Although transfer learning improved (smaller KSD values against native) the generation quality of weaker models in all 10 antigens for the within antigen transfer scenario, three antigens (3RAJ, 3VRL, and 5E94) did not show improvements (larger KSD values against native) for the across antigens transfer scenario (although closer examination of the generated affinity distributions revealed that the median affinity values of across antigens transfer learning were closer to the median affinity values of native CDR-H3 sequences, Fig. S6). Furthermore, the number of recognized epitopes in any transfer learning was notably larger than the number of recognized epitopes in sequence generation without transfer learning and in native CDR-H3 sequences (Fig. S13) independent of the KSD values against native CDR-H3 sequences. This illustrates the key challenges remaining in the prospective testing of many orthogonal variables wherein several parameters must be captured and justly reflected in order to communicate faithfully the underlying trends in the data. Indeed the success of cross tasks transfer has been shown to be heavily influenced by the compatibility of the source and target task types (*40*). Nevertheless, our cross-antigens transfer learning experiments show that, at least in the case of our antibody sequence datasets, neural network models can extrapolate 3D non-linear dependencies to CDR-H3 sequences outside the training distribution (*40–42*).

One may argue that our framework generates sequences that are binders within the lattice framework but would not be binders if tested in vitro/vivo. That said, we ensured that the antibody-antigen simulation framework is state-of-the-art surpassing all currently available large-scale antibody-antigen binding simulation frameworks (*28*) (e.g., the inclusion of discretized PDB-stored antigens, 3D-binding [albeit on a 90°-grid], experimentally determined physiologically relevant amino-acid interaction potentials (*26*)). The inbuilt physiological relevance of our antibody-antigen simulation model affords a more precise understanding of how the accuracy of computational models increases with the number of available antibody sequences for training, which will help in planning experimental validation studies. We also avoided the possibility that the generative model learns to exploit the affinity models by refraining from a full reinforcement learning setting, in which the affinity model would be used as a reward function (Renz et al., 2020). Specifically, the major challenge of predicting antigen reactivity of an antibody sequence lies in recapitulating the residue interactions between the antibody and antigen structures in 3D space. Even our simplified computational model of antibody structure includes physical antibody-antigen interactions in 3D space entailing non-linearities and positional dependencies reminiscent of the biological complexity (*26*). Consequently, one may argue that our simulation framework and investigations are suitable for establishing an informative lower bound of the complexities encountered in machine and deep-learning-based biological sequence design. Indeed, a recent study by Mason et al. (*31*) that leverages experimental deep mutational scanning data showed that a training dataset size in the order of 10^4^ (as also shown in this study; Figure 4A) was sufficient to train machine learning models that discriminate binders and non-binders. Furthermore, the study also highlights that a large proportion of dissimilar sequences (LD>2) bind to the target antigen (as also shown in this study; Figure 2D). These parallels (with results from experimental data) reiterate the utility and relevance of simulated custom-designed synthetic datasets in advancing the development of computational approaches for antibody design and discovery.

For future investigations, we cannot highlight enough the need for experimental validations to compliment the herein *in silico* results. Recently, Saka and colleagues showed that RNN-based generated antibody sequences bind the desired target providing experimental proof of principle of our computational framework (*43*). Here, we validated our RNN-LSTM framework by scoring the generated CDR-H3 sequences using an experimentally validated oracle (CNN-classifier) (*31*) (Figure 5). Furthermore, our RNN-LSTM models were trained separately on binders (positive data) and non-binders (negative data) suggesting that the design of native-like CDR-H3 sequences is possible without the need for negative examples and accuracy is likely to be further improved with more training examples (the 72%-HER2 generation rate by the RNN is fairly close to the CNN prediction accuracy of ≈80%, (*31*)). This could potentially reduce the cost to generate training datasets given that the HER-2 generation rate of the RNN was remarkably high despite only being trained on positive data. Indeed earlier studies have shown that performance improvements/reductions by including more or less negative data vary across models and application domains (*44, 45*). This highlights the potential applicability of our framework in real-life settings beyond the synthetic simulated setting earlier described. We strongly believe that the synergistic combination of simulation and experimental strategies is necessary for the time- and cost-efficient discovery of antibody therapeutics. Naturally, future refinements to the Absolut! simulation framework would further improve the applicability of conclusions drawn to experimental settings. These refinements are among others (see (*26*) for a more detailed discussion): (i) antibody full VH-VL chains (so far, we can only model CDRH3-antigen binding), (ii) smaller angle grid in the lattice: our framework was limited to integer positions in a 3D grid, (iii) addition of constraints at the CDR3 ends in order to reproduce the anchoring of the CDR chains to the framework/conserved domains of the antibody.

Once more experimental data have become available, one may venture into merging simulation and experimental training data. For example, one could perform transfer learning based on antibody sequences with only partially determined experimental labels thus increasing the biological faithfulness of deep-learning-designed antibody sequences (*46*). Such a setup may be further augmented in the form of federated learning (*47*). Furthermore, here we performed deep learning on amino acid sequences and not nucleotide sequences although nucleotide sequences are essential for experimental antibody expression. However, codon usage is often species-specific (*48*). Therefore, we opted for the more general amino acid encoding. Nevertheless, our deep learning setup would work equally well for nucleotide sequences.

A key property of *in silico* generative frameworks such as ours is that once trained, it paves the way for large-scale and on-demand generation of antigen-specific and developable immune receptor sequences. The fast production of antibodies has seen continued interest from the field (*6, 7*). Although library-based discoveries have the potential to generate a higher volume of antigen-specific data as compared to crystallography or related approaches, they remain reliant on multiple rounds of selection as well as other experimental heuristics. We approached the discovery process by leveraging deep generative models, which implicitly aim to learn the rules of antibody-antigen binding. Once learned, the generation of vast quantities (virtually limitless) of antibody sequences becomes feasible, abrogating the need for follow-up screening. Rule-based generation also imparts the ability to design (not merely discover) antibody sequences by biasing the deep generative models towards a particular set of developability parameters via reinforcement learning or instance selection (*18, 49*). The combination of near-limitless and fast sequence generation may enable the construction of an on-demand antibody generator where antigen-specific antibody sequences can be obtained at will.

In this work, we did not train on datasets that were selected for both binding and developability therefore not optimizing both antibody design entities at once. This is partly due to the inherent sparsity of the data although our datasets are the largest currently available. Incorporating several orthogonal properties in one training dataset is an interesting avenue for future research. Furthermore, we would like to point out that we have not optimized in any way the deep generative architecture used. Therefore, our framework allows for optimizing the generative output of deep learning approaches in future benchmarking studies (*10, 50*).

In closing, naturally occurring proteins represent only a small subset of the theoretically possible protein sequence space. Here, we demonstrate a proof-of-principle that deep learning helps explore a broader sequence and structural space than present in the training data thereby enabling the discovery and the design of antibody sequences with enhanced or novel properties (*6, 51*). Moreover, our ground-truth-based framework may be useful in the establishment of methods for model interpretability (*51–54*).

## Methods

### Reference experimental B-cell receptor and 3D-crystal structure antigen data

Native B-cell receptor (CDR-H3) sequences (*n*_seq_ = 7 × 10^6^, murine origin [we showed in a separate work that murine and human CDR-H3 sequences have similar affinity distributions in the Absolut! antibody-antigen simulation framework] (*26*)) were obtained from Greiff and colleagues (*33*). Ten antigen 3D-crystal structures were sourced from known antibody-antigen complexes in the Antibody Database (AbDb) (Table 1) (*32*) (Table 1) and converted into lattice-based discretized Absolut! format (*26*). To annotate each CDR-H3 sequence for antigen specificity, we determined the best binding position of an antibody sequence to an antigen and calculated the corresponding binding affinity via the software suite Absolut! (see below and (*26*)). Antigen-specific CDR-H3 sequences were defined as the top 1% affinity-sorted CDR-H3 sequences for each antigen (n_seq_=1% times 7×10^6^= 70 000). We chose the top 1% as it selected a sufficiently high number of sequences as well as ensured high antigen-specific affinity (see Supplementary Fig. S3 for a comparison of the distribution of all 7 million CDR-H3 sequences [“native”] vs the top 1% affinity ones [“native_top”]).

### Reference CNN model trained on experimental human epidermal growth factor 2 (HER2) CDR-H3 binder and non-binder sequences

CDR-H3 sequences that bind (binders) and do not bind (non-binders) to HER2 were obtained from Mason and colleagues (*31*). They used a total of experimentally validated 11 300 HER2-binders and 27 539 validated HER-2 non-binders to train a convolutional neural network (CNN) classifier that assigns an HER-2 binding probability to a given input CDR-H3 sequence. The accuracy of this CNN classifier was experimentally validated. We used the CNN classifier to evaluate the HER-2 binding probability of CDR-H3 sequences generated by our RNN-LSTM model (see Figure 5).

### Generation of lattice-based antibody-antigen binding structures using Absolut!

The Absolut! software was used to compute the binding energy and best binding structure (here termed paratope fold or binding fold) of antibody (CDR-H3) sequences around the antigens in a 3D-lattice space (see (*26*) for a very detailed explanation). Briefly, the antigens of interest, named by their PDB entry (Table 1), were transformed into a coarse-grained lattice antigen representation (a step called discretization, performed using the program LatFit (*55*)]), where each residue occupies one position and consecutive AAs are neighbors, creating a non-overlapping 3D chain with only 90 degrees angles. In the lattice, a position is encoded as an integer (for instance, [x=31, y=28, z=15] is encoded as a single integer code [x+L*y+L*L*z] where L is the lattice dimension: 64). Further, protein chains are represented as a starting position and a list of moves, for instance, 63263-SUSDLLUR is a peptide starting at position (x=31, y=28, z=15; 31+64*28+64*64*15=63263) with 9 AAs and following the structure ‘Straight, Up, Straight, Down, Left, Left, Up, Right’ where each ‘turn’ is defined from the previous bond and is coordinate-independent. From each CDR-H3 sequence investigated, all peptides of 11 consecutive AAs are taken (sliding window with a step size of 1; window size of 11 was chosen to provide the best compromise between computational cost and CDR-H3 length/coverage see (*26*)) and are assessed for binding to the antigen. From exhaustive enumeration of all possible structures of the peptide around the antigen, Absolut! returns the structure minimizing the energy of the complex (Fig. S1). Exhaustive enumeration of all possible binding folds (binding structures) of a CDR-H3 sequence enables Absolut! to function as an oracle since it can generate the binding fold as well as evaluate the binding energy of any sequence against the antigen of interest. The energy is computed from neighboring, noncovalent AAs either between the CDR-H3 and the antigen (binding energy) and within the CDR-H3 (folding energy) using an empirical experimentally estimated potential (*56*). Among all 11 AAs peptides for this CDR-H3, the one with the best total (binding + folding) energy is kept and its structure is called the ‘binding structure’ or the ‘paratope fold’ of the CDR-H3 (Fig. S1). The paratope in that structure will be the spatial conformation of interacting AAs on the antibody side and the epitope the spatial conformation of interacting AAs on the antigen side. In that way, each CDR-H3 sequence is annotated with a 3D binding structure (paratope and epitope) and binding energy (see Figure 1 and Fig. S1 for illustration). In summary, using Absolut!, we constructed a dataset of 70 million (7 Million CDR-H3 sequences x 10 antigens [Table 1]) antibody-antigen structures with annotated paratope, epitope, affinity, and antibody developability (see below).

### Computation of developability parameters

Developability is defined as the “feasibility of molecules to successfully progress from discovery to development via evaluation of their physicochemical properties” (*57*). Developability parameters (Table 2, inspired by the works described in (*18, 31, 58*) were computed using the module `Bio.SeqUtils.ProtParam` in `Biopython` (*59*) and `NetMHCIIpan` versions 4.0 and 4.1 (*60*). For `NetMHCpan` and `NetMHCIIpan` we used the percent rank (the percentile of the predicted binding affinity compared to the distribution of affinities calculated on a set of random natural peptides) where typically the thresholds for strong binders are defined at 2% and weak binders between 2-10%.

### Deep generative learning using long short-term memory neural networks

The architecture of the deep generative model used consists of three layers (see Figure 1 and Fig. S2): (i) an embedding layer with 256 output-vector dimensions, (ii) a recurrent neural network of the type long short-term memory (RNN-LSTM) (*61*) with 1024 units and finally (iii) a fully-connected output layer with softmax-activations of 21 (twenty amino acids and one whitespace character) output-vector dimensions (see Figure 1). Input-target pairs, i.e., sequences and their labels, were obtained by first merging the antibody sequences (CDR-H3s) into a text corpus, sequences were separated by a single whitespace character. A window of size *w* (*w*=42 amino acids) was used to fragment the corpus into chunks of input sequences *x* of length *w*. For each input sequence *x* a target sequence *y* was created by sliding a window of size *w-1* one step forward. The last character was removed from *x* creating an input-target pair (*x,y*) each with the size *w-1*. Thus, the LSTM model *g*(*x; θ*) is trained to predict the next character of the given sequence using categorical cross-entropy loss *L*(*y*, *g*(*x, θ*)), where *θ* is the parameter/weight of the LSTM model. We partitioned the input-target pairs into training (70%), validation (15%), and test (15%) sets. The training was carried out for 20 epochs with Adam optimizer (*62*). At the end of each epoch, training and evaluation loss were computed for evaluation. The generation was initiated with a seed string and the hyperparameter temperature was set to 1. Our implementation is based on TensorFlow 2.0 (*63*).

### Implementation of transfer learning

We leveraged transfer learning to examine whether the generation quality of models trained on lower-sized datasets (data-poor, “low-N, (37”) may be improved by transferring learned features from models trained on larger training datasets. For a visualization of the transfer learning setup, see Figure 4. Prior to any transfer learning experiment, we randomly sampled 1% (n_sample_=700) and 10% (7 000) sequences from the set of antigen-specific CDR-H3 sequences, defined as the top 1% affinity-sorted CDR-H3 sequences for each antigen, n_sample_=70 000). Sampling was performed 5 times per antigen. Models trained on 70 000 sequences were termed “data-rich” and models trained on 700-7000 sequences, “data-poor”. In a transfer experiment, a *transfer learning architecture* was constructed by stacking the pre-trained embedding followed by the pre-trained RNN-LSTM layers from data-rich models and a new fully-connected layer (see Figure 4B for network architecture). The training was performed as described in the previous section (“Deep generative learning”). Transfer learning was performed in two ways termed “within-antigen” and “across-antigens” (see Figure 4). “Within-antigen” experiments describe transfer-learning between the same antigen (e.g., embedding and RNN-LSTM layers of data-rich models stem from the same antigen that is used to train a data-poor model). “Across-antigens” describes the transfer of layers between different antigens (e.g., the combination of a data-rich model of antigen V and a data-poor model of antigen G). The within-antigen experiments served as positive controls where stronger signals (from a data-rich model) were used to improve the performance of a weaker model.

### Sequence similarity, composition, and long-range dependencies

Sequence similarity among generated CDR-H3 sequences was determined by Levenshtein distance and gapped k-mer analysis. Levenshtein distances were computed using the distance function in the package `Python-Levenshtein` (*64*). Long-range dependencies were assessed by gapped k-mer analysis using the R package kebabs (*65*) as previously described (*66, 67*).

### Distance between distributions

The similarity between two CDR-H3 affinity distributions was quantified using the Kolmogorov-Smirnov distance (KSD) using the function `ks.diss` (Kolgomorov-Smirnov test, KSD) from the R package `Provenance` (*68*). The KSD measures the largest vertical distance between the two examined (cumulative) distributions. A KSD value close to 0 indicates that the distributions are very similar and a larger distance (e.g., 1) indicates larger differences between the distributions.

### Mean squared error of positional amino acid frequency matrix

As previously described (*69*), the difference between two amino acid position-specific frequency matrices (Figure 5) was quantified by the mean squared error (MSE) 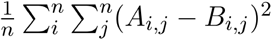, where *A* is the reference native amino acid frequency matrix, *B* is the generated amino acid frequency matrix, *n* is the twenty amino acid alphabet, *m* is the length of CDR-H3, *i* is the row index and *j* is the column index.

### Graphics

Plots were generated using the R package ggplot2 (*70*) and arranged using Adobe `Illustrator 2020` (Adobe Creative Cloud 5.2.1.441).

### Hardware

Computations were performed on the Norwegian e-infrastructure for Research & Education (NIRD/FRAM; https://www.sigma2.no) and a custom server.

### Data and code availability

Preprocessed datasets, code, and results figures are available at: https://github.com/cs?-greifflab/manuscript_insilico_antibody_generation.

## Acknowledgments

We acknowledge generous support by The Leona M. and Harry B. Helmsley Charitable Trust (#2019PG-T1D011, to VG), UiO World-Leading Research Community (to VG), UiO:LifeScience Convergence Environment Immunolingo (to VG, GKS, and IHH), EU Horizon 2020 iReceptorplus (#825821) (to VG), a Research Council of Norway FRIPRO project (#300740, to VG), a Research Council of Norway IKTPLUSS project (#311341, to VG and GKS), a Norwegian Cancer Society Grant (#215817, to VG), and Stiftelsen Kristian Gerhard Jebsen (K.G. Jebsen Coeliac Disease Research Centre) (to LMS and GKS). This work was carried out on Immunohub eInfrastructure funded by University of Oslo and jointly operated by GreiffLab and SandveLab (the authors) in close collaboration with the University Center for Information Technology, University of Oslo, IT-Department (USIT). Finally, we thank Derek Mason and Sai T. Reddy for constructive comments and discussion.

## Supplementary Material

**Supplementary Figure S1.**
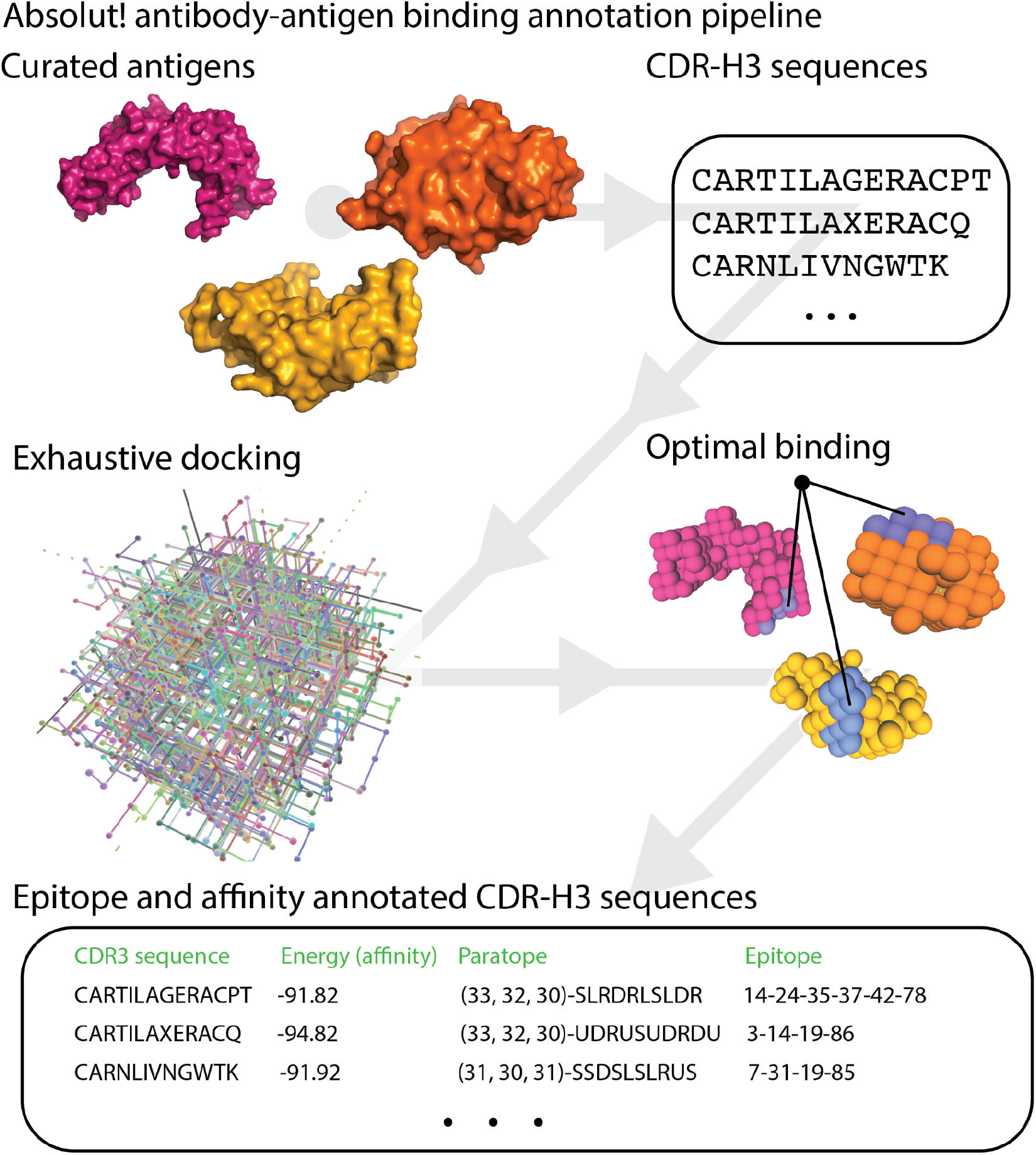
(relates to Figure 1 and Figure 2B). Absolut! Antibody-antigen binding pipeline (*26*). For a given antigen and CDR-H3 sequence, the software suite Absolut! performs an exhaustive search (termed exhaustive docking (*26*)) over all binding positions, returns the corresponding binding affinities, and finally outputs the binding energy (affinity) and the binding structure (the paratope fold, paratope amino acid residues, and epitope amino acid residue position) for the optimal (lowest energy) binding. A protein chain is defined by its amino acid sequence, a starting point in space, and a list of relative moves in space to determine the next AA position (Straight (S), Up (U), Down (D), Left (L), Right (R)).

**Supplementary Figure S2.**
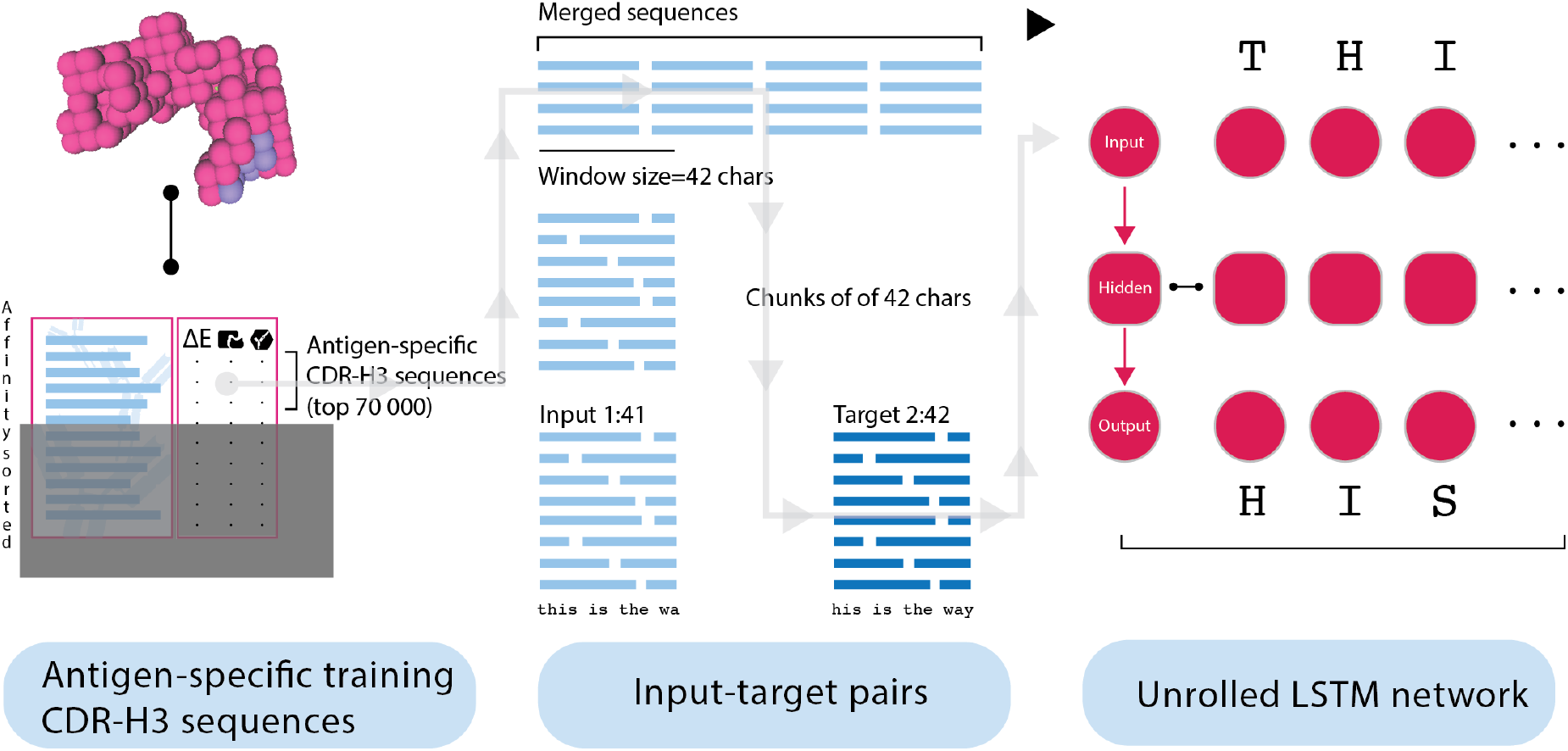
(relates to Figure 1). Workflow of deep generative learning using long short-term memory neural networks. To create input-target pairs to train the network (i) we defined the top 1% of affinity sorted CDR-H3 (amino acid) sequences (n_seq_=70 000) as antigen-specific sequences, (ii) merged the sequences into a text corpus, (iii) fragmented the text corpus into segments sequences of size *w*=42, and (iv) created input-target pair of 1:w-1 and 2:w characters. The input-target pairs are then used to train the LSTM network.

**Supplementary Figure S3.**
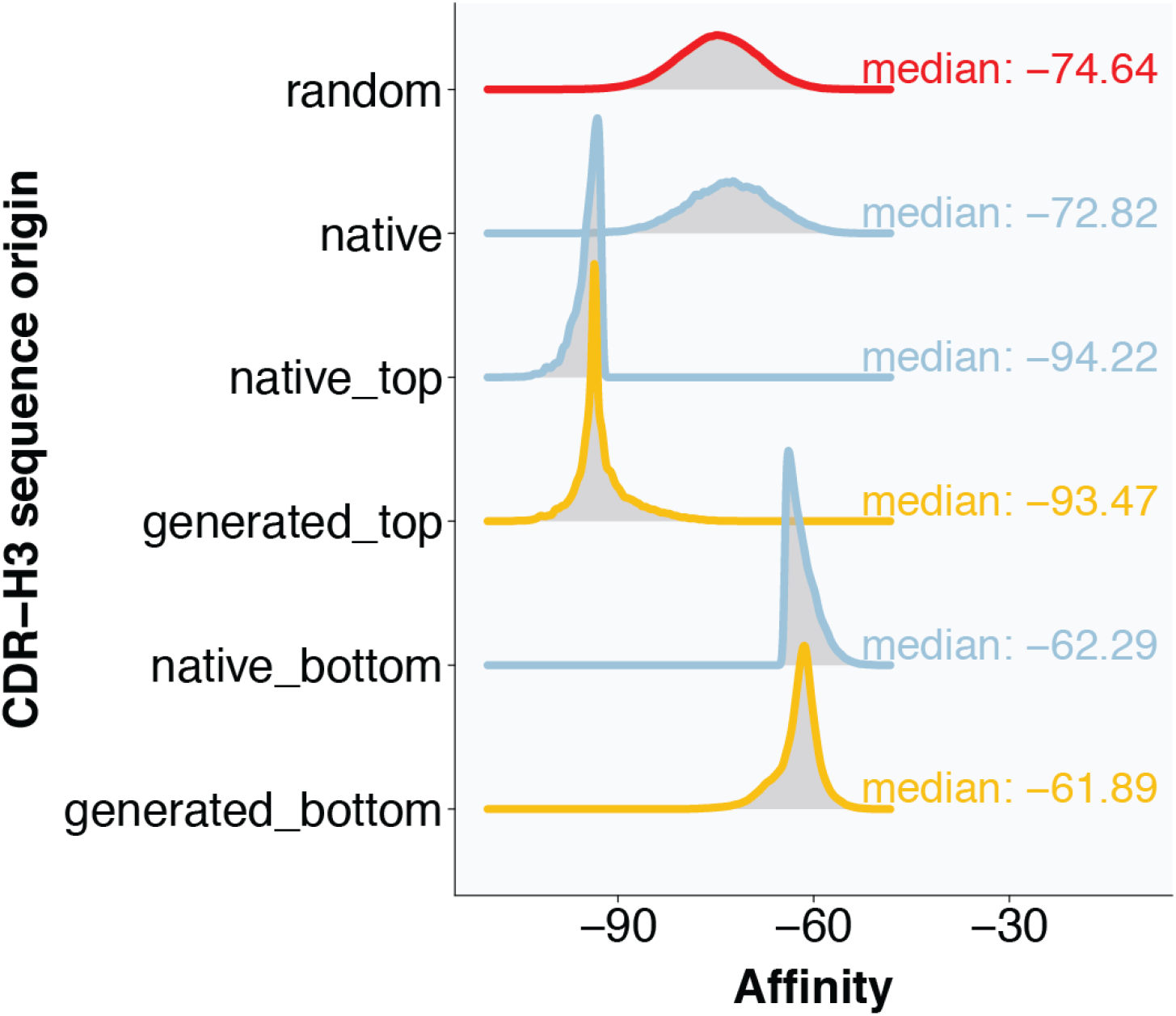
(relates to Figure 2A). Generated CDR-H3 sequences of models trained on high and low-affinity sequences reproduce the high and low binding affinity distribution accordingly. To examine the capacity of our model to generate sequences that are localized at different regions of the native binding affinity spectrum, we trained the model, separately, on high and low-affinity sequences (high and low-affinity sequences were obtained by first sorting the native sequences according to their binding affinity and subsequently taking the top and bottom 1% of the affinity-sorted data for the antigen 1OB1; n_seq_top_: 70 000; n_seq_bottom_: 70 000). We found that generated sequences sufficiently reproduce the binding affinity distributions of top and bottom sequences. generated_top_median_: −93.47, yellow; native_top_median_: −94.22, blue; generated_bottom_median_: −61.89, yellow; native_bottom_median_: −62.29, blue. As a baseline, we also displayed the binding affinity distribution of all 7×10^6^ native sequences, median: −72.82, blue; and the binding affinity distribution of random (sampled from a uniform amino acid distribution, n_seq, random_=10^6^) sequences, median: −74.64, red.

**Supplementary Figure S4.**
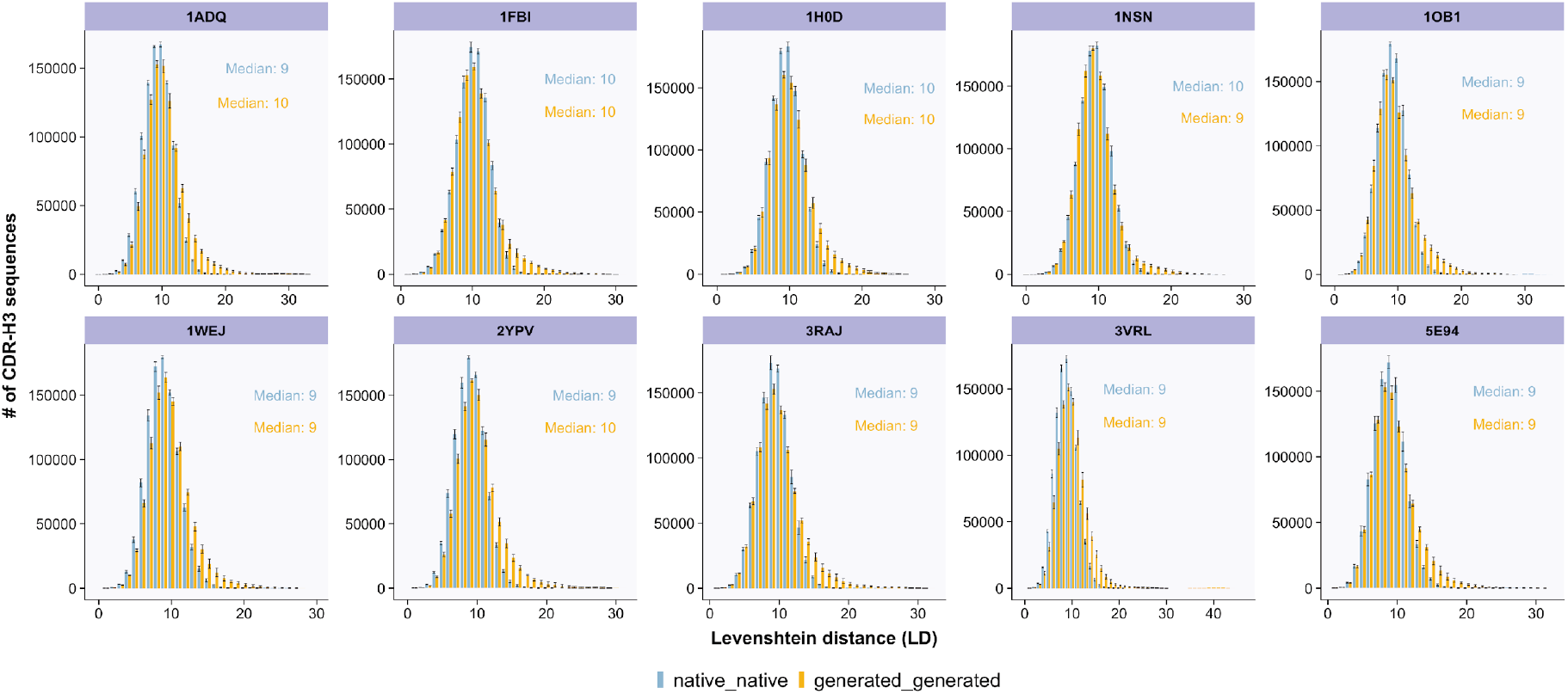
(Relates to Figure 2D) Distributions of Levenshtein distances calculated among native and generated CDR-H3 sequences are largely similar. To quantify how similar (or different) native and generated CDR-H3 sequences are, we calculated Levenshtein distances (LD) for native and generated CDR-H3 sequences of a given antigen and plotted the distances as distributions. Medians of each distribution are displayed. We found that the native and generated distributions overlap substantially. (for a more tractable pairwise LD calculation, we subsampled the full dataset five times (n_seq,subsample_=1000), showed the mean of the number of sequences per LD, and used the resulting standard deviations as error bars.)

**Supplementary Figure S5.**
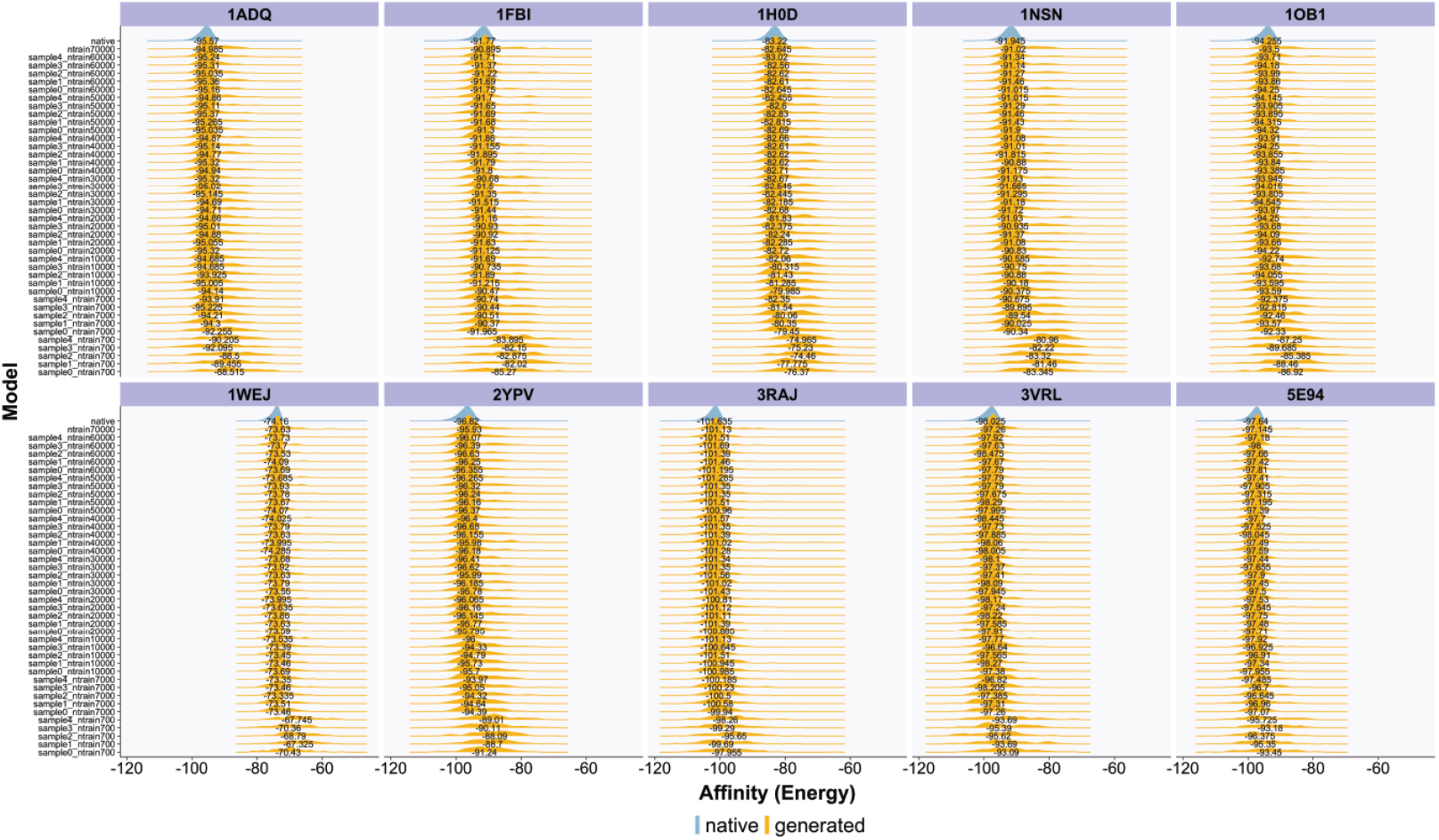
(relates to Figure 4A) Affinity (as indicated by energy) improves as a function of the number of training sequences. To examine the impact of sample size on the resulting binding affinity of generated CDR-H3 sequences, we created smaller training datasets (n_seq,subsample_=700, 7000, 10 000, 20 000, 30 000, 40 000, 50 000, 60 000, and n_replicates_=5) from the full antigen-specific CDR-H3 sequences (n_seq,training_=70 000), trained deep generative models on the subsets and compared the binding affinity and epitope against affinity and epitope from models trained on the full data and the native affinity and epitope. We found that models trained on the larger dataset sizes (>2×10^4^), but not the smaller subsets (in the order of 10^3^ or 10^2^), sufficiently replicate the distribution of binding affinity and epitope CDR-H3 sequences. Medians of these distributions are shown in Figure 4A as boxplots.

**Supplementary Figure S6.**
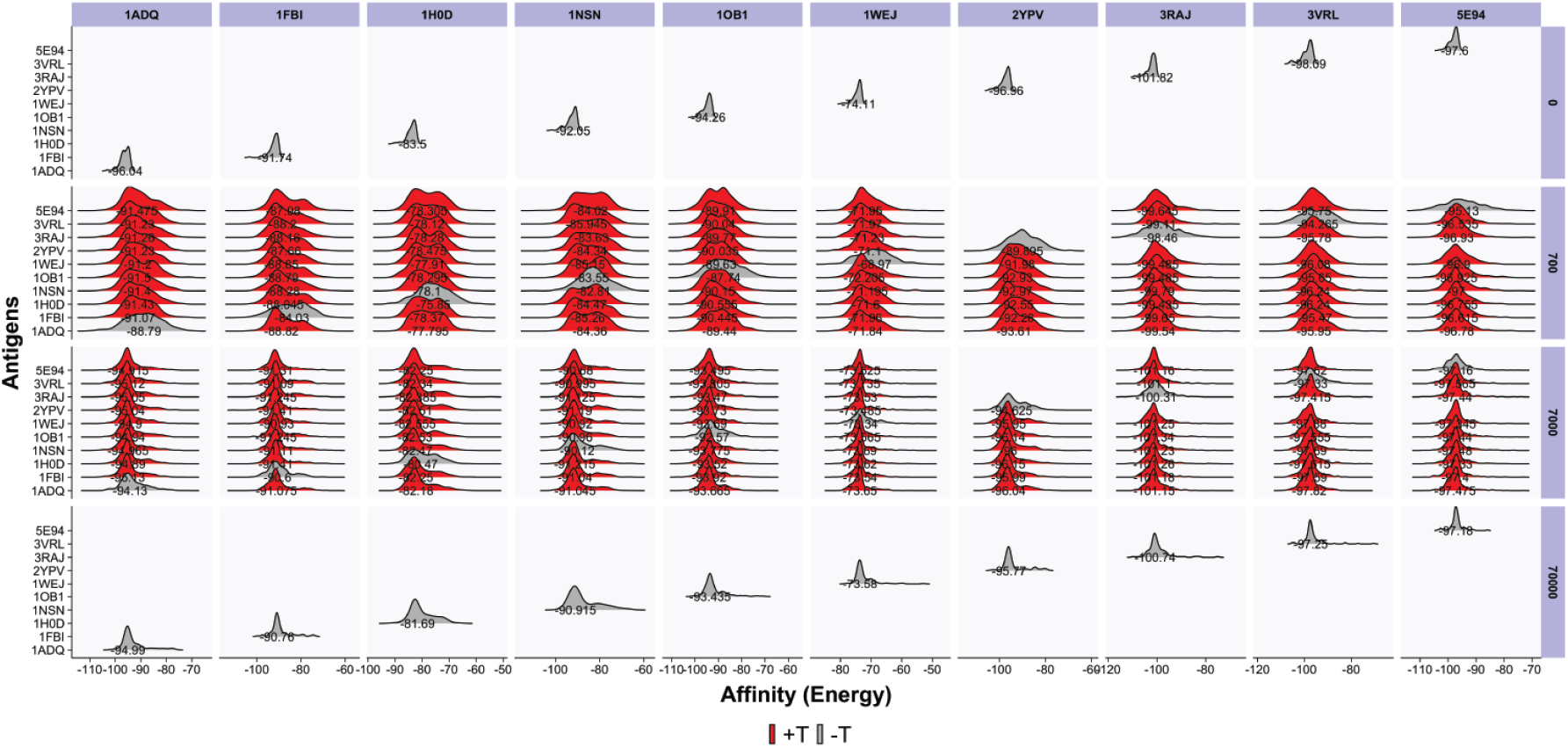
(relates to Figure 4B) Distributions of cross-transferred binding affinities. To examine the impact of cross-transfer learning (transfer learning across antigens), we (re)used the embedding and LSTM layers of data-rich models (N_seq, training_=70 000, row: 4) in combination with a fresh dense layer, train the resulting model with reduced training datasets (N_seq, training_: 700/7000, row: 2 and 3), and compared the binding affinity of the generated sequences against the native (N_seq, training_: 0, row: 1) and data-rich (N_seq, training_: 70 000, row: 4) affinities (see Figure 4). Donor and recipient antigens are denoted in the columns and rows (of each panel), respectively. For purposes of readability, here shown are aggregate distributions (the composite of the five random samplings for each cross-transfer task). Broadly, we found that the medians of affinity distributions of the cross-transferred models (red, row 2 and 3) were closer to the medians of affinities of the native and data-rich distributions (grey, row 1 and 4) compared to the medians of models without transfer (grey, row 2 and 3).

**Supplementary Figure S7.**
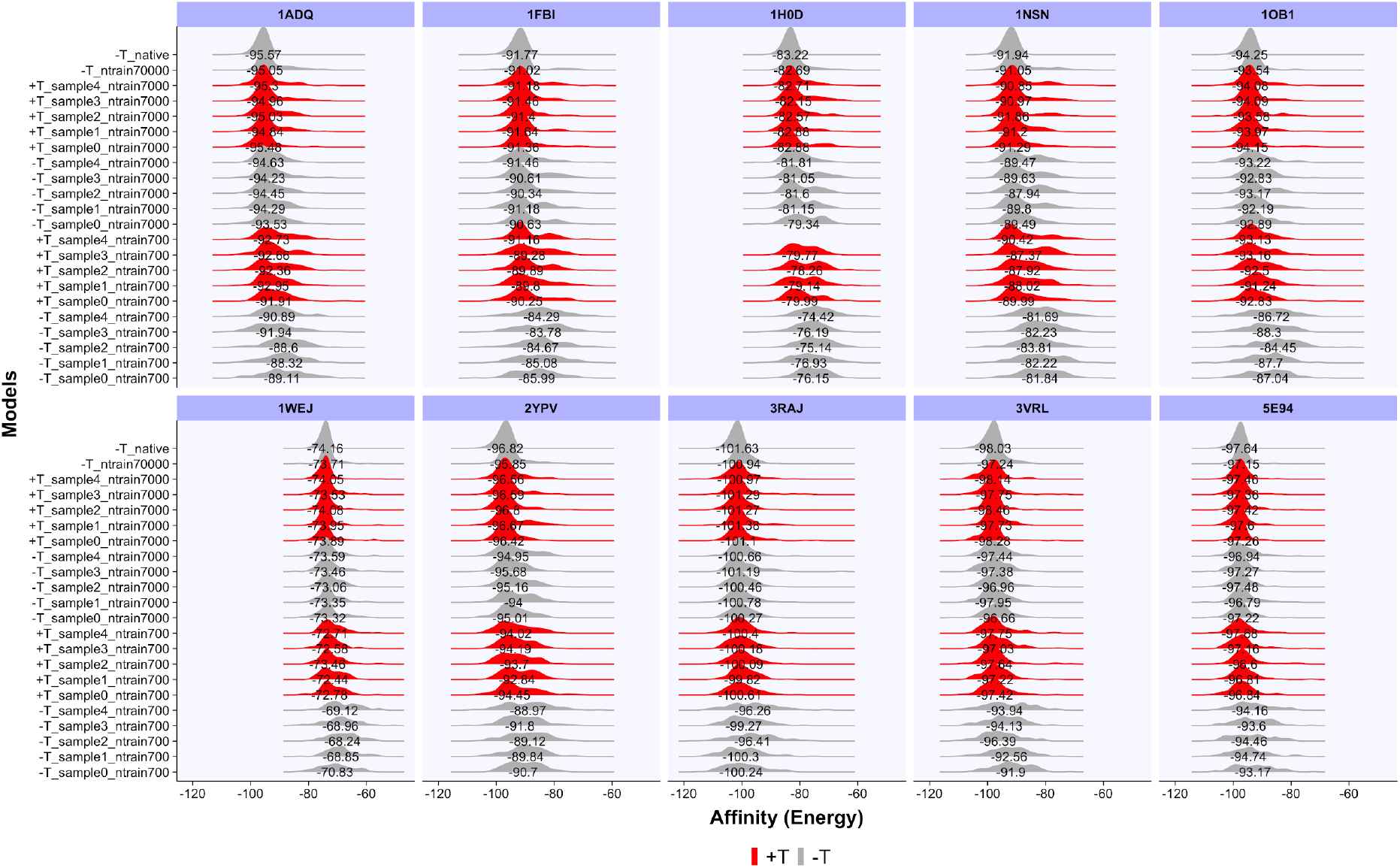
(relates to Figure 4B) Distributions of transferred affinities. Transfer learning was performed by (i) constructing a transfer architecture wherein embedding and RNN-LSTM layers from a “data-rich” model (n_seq, training_=70 000) were stacked atop of a fresh dense layer and (ii) training the resulting ‘transfer’ model on lower-sized datasets (data-poor, n_seq, training_=700, 7000). Within-antigen transfer experiment describes transfer learning within the same antigen (e.g., between a data-rich model of an antigen *V* and data-poor models of the same antigen *V*). Affinity distributions of CDR-H3 sequences generated by transfer (+T/with transfer) models are shown in red whereas affinity distributions of CDR-H3 sequences generated by non-transfer models (-T/without transfer) are shown in grey. Medians of these distributions are shown in Figure 4B as boxplots.

**Supplementary Figure S8.**
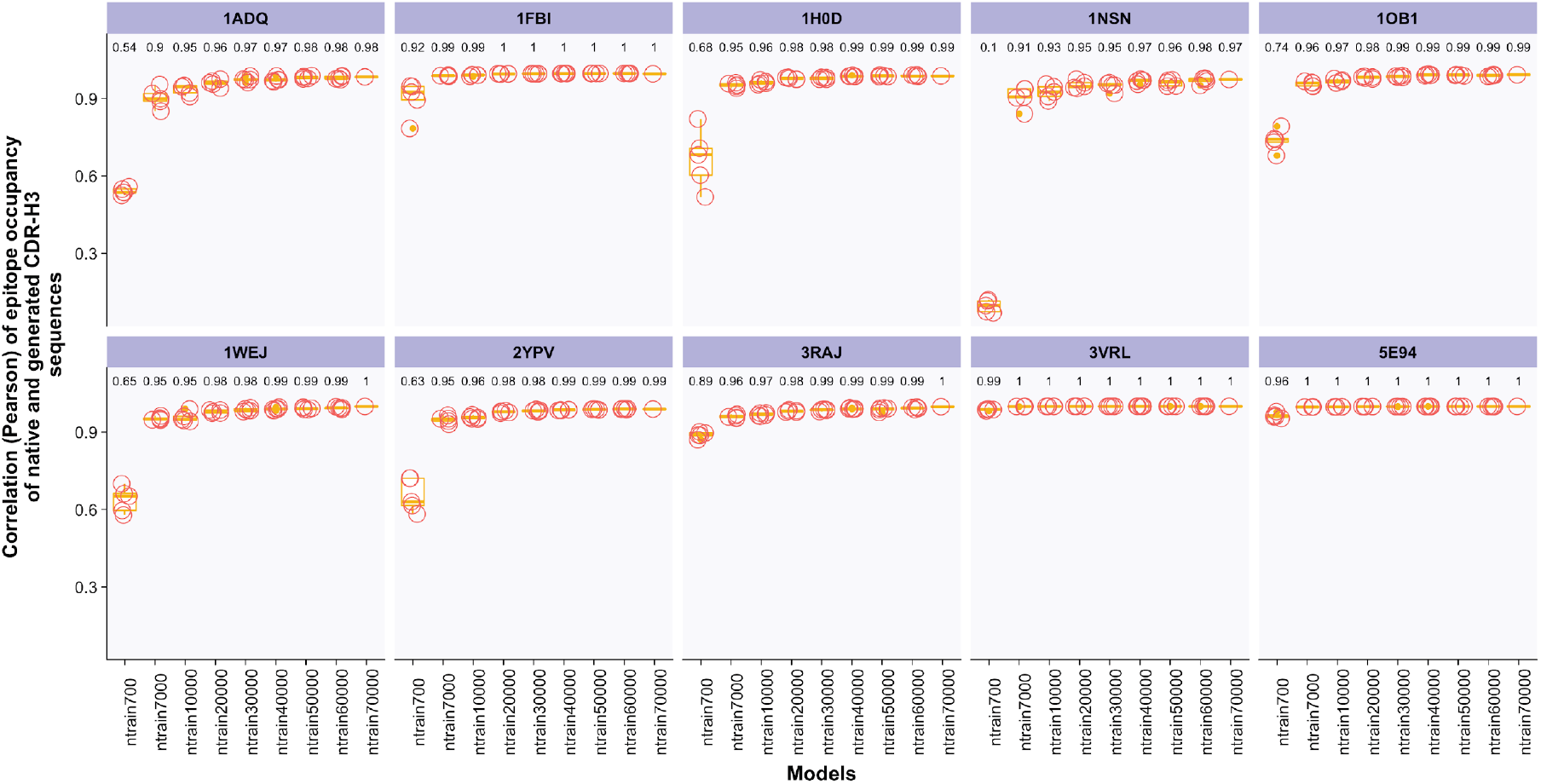
(relates to Figure 4A) Distributions of Pearson correlation of frequencies of epitopes recognized by native and generated CDR-H3 sequences as a function of the number of training sequences. To examine whether generated CDR-H3 sequences recovered the epitopes of the native CDR-H3 sequences, we correlated the frequencies of epitopes recognized by native and generated CDR-H3 sequences for all antigens (n_sample_=5 per model). Broadly, we found that the Pearson correlation values increase as a function of the number of training sequences. In addition, we found that the Pearson correlation for antigens with fewer epitopes (e.g., 3VRL, Figure 2B bottom panel) was already reasonable at a small training dataset (ntrain700). In contrast, the correlation for antigens with many epitopes (e.g., 1H0D, Figure 2B, bottom panel) required larger training datasets for reaching a high concordance with the epitope occupancy found in the training dataset.

**Supplementary Figure S9.**
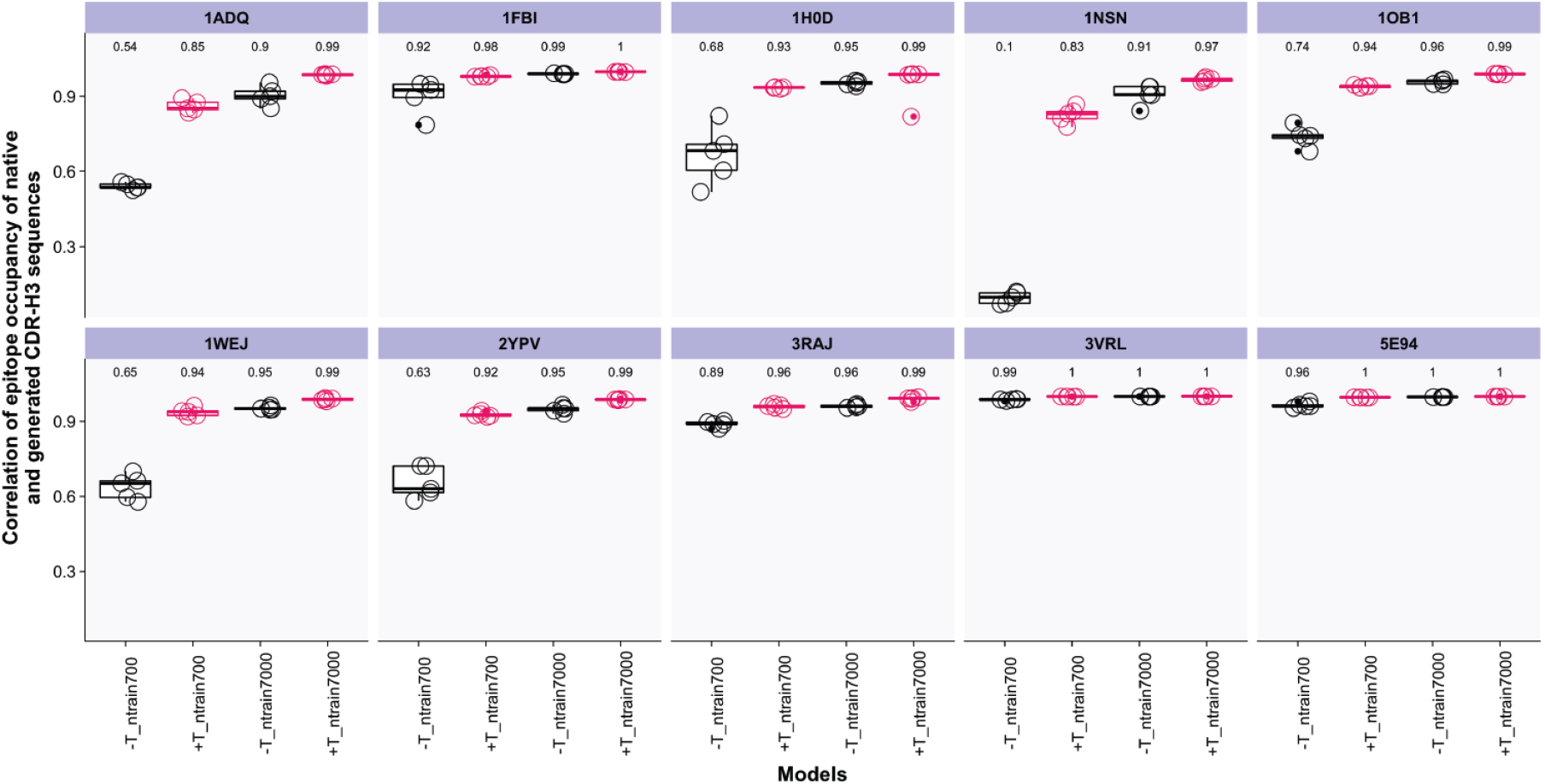
(relates to Figure 4B, within antigen transfer) Distributions of Pearson correlation of frequencies of epitopes recognized by native and generated CDR-H3 for the within antigen transfer setting. To examine whether within-antigen-transfer-generated CDR-H3 sequences recovered the epitopes of the native CDR-H3 sequences, we correlated the frequencies of CDR-H3 bound to epitopes bound by native and transfer-generated CDR-H3 sequences for all antigens for small dataset sizes as described in Figure 4B. We found that epitope occupancy was higher correlated for CDR-H3 sequences with transfer learning (+T) compared to those generated without transfer learning (-T) in particular for the smallest dataset size (n_seq, training_ = 700).

**Supplementary Figure S10.**
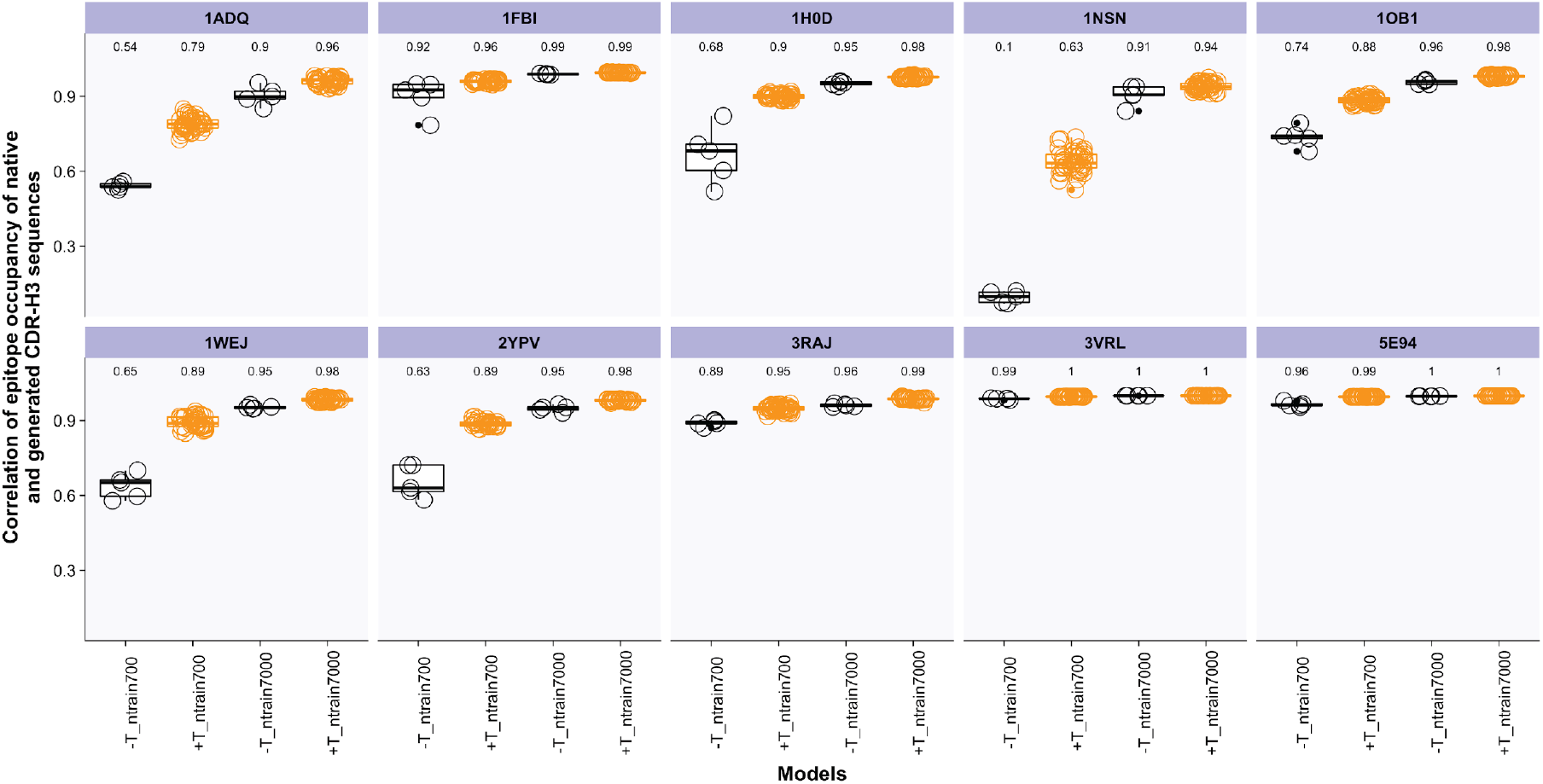
(relates to Figure 4B, across antigens transfer) Distributions of Pearson correlation between CDR-H3 frequencies bound to native and transfer-generated epitopes in the across-antigens setting. To examine whether across-antigens-transfer-generated CDR-H3 sequences recovered the occupancy of the epitopes of the native CDR-H3 sequences, we correlated the frequencies of CDR-H3 bound to epitopes bound by native and transfer-generated CDR-H3 sequences for all antigens for small dataset sizes as described in Figure 4B. We found that epitopes were better recovered in CDR-H3 sequences with transfer learning (+T) compared to without transfer learning (-T) in particular for the smallest dataset size (n_seq, training_ = 700).

**Supplementary Figure S11.**
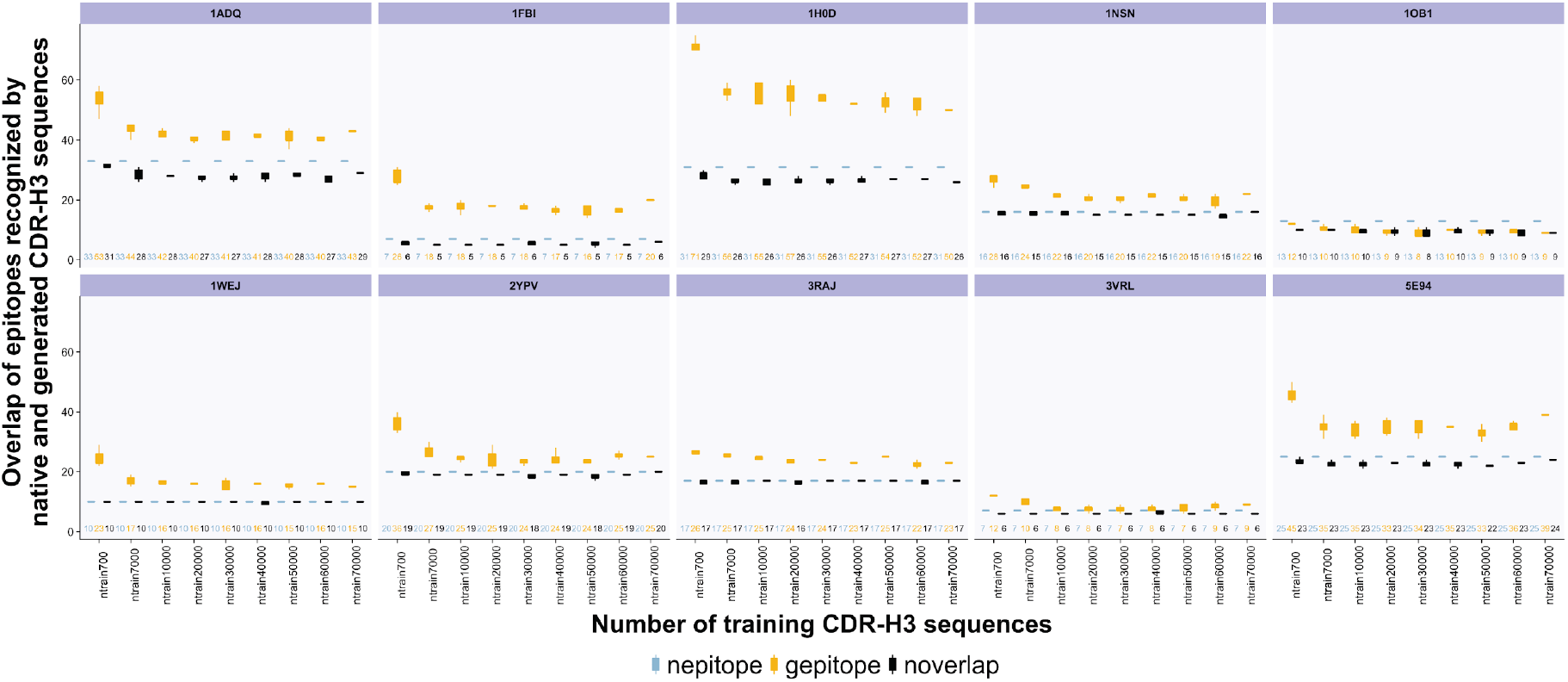
(relates to Figure 4A and Fig. S8) Overlap of epitopes recognized by native and generated CDR-H3 sequences as a function of the number of training CDR-H3 sequences. For each training dataset size (ntrain700-ntrain70000), we plotted the total number of epitopes recognized by native (training) CDR-H3 sequences (*nepitope*), the total number of epitopes recognized by generated epitopes (*gepitope*), and the absolute overlap of epitopes recognized by native and generated CDR-H3 sequences. (*noverlap*). Broadly, we found that generated CDR-H3 sequences recognized a higher number of epitopes (except for 1OB1) than native CDR-H3 sequences.

**Supplementary Figure S12.**
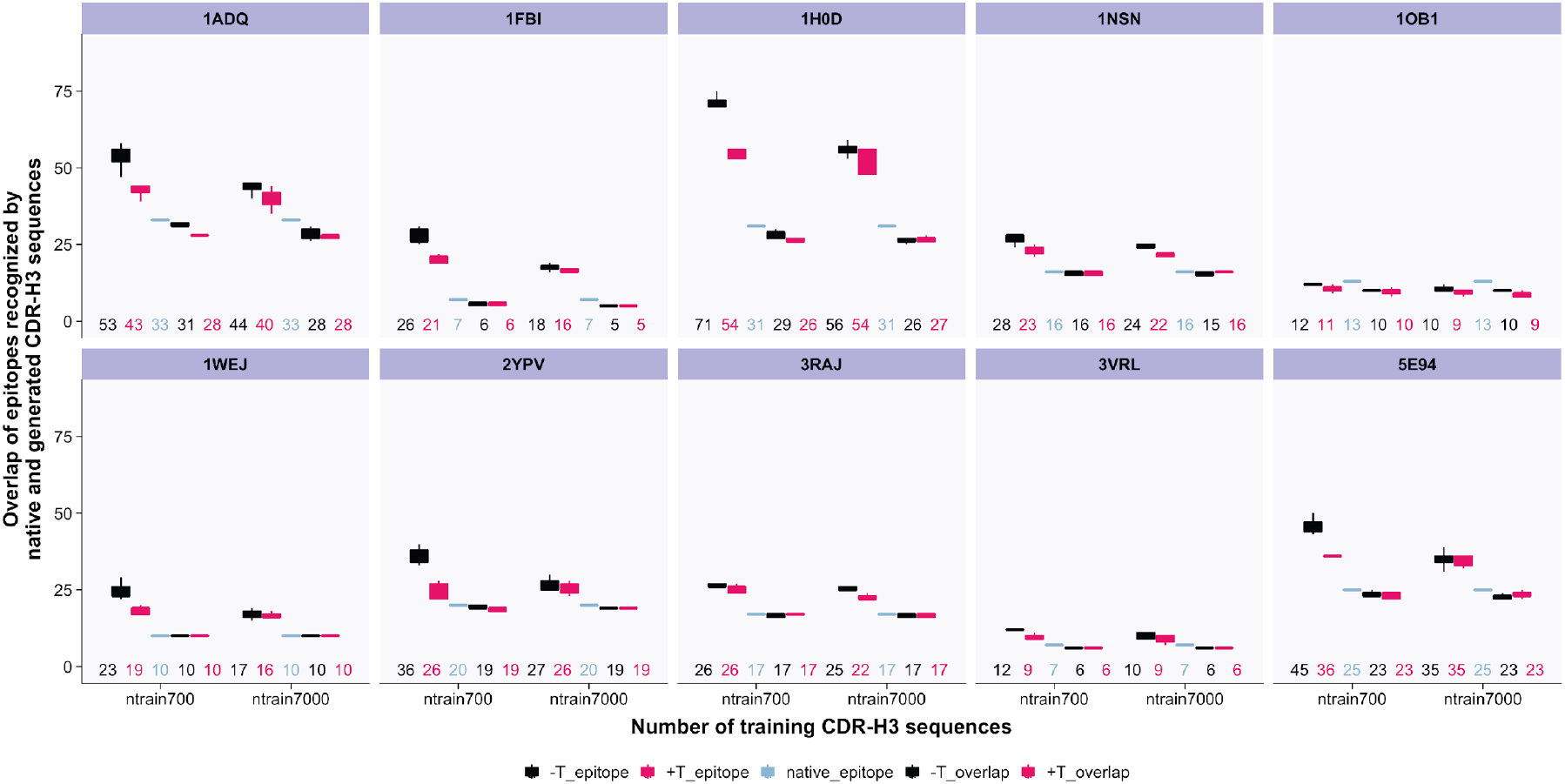
(relates to Figure 4B top panel and Fig. S9). Overlap of epitopes recognized by native and generated CDR-H3 sequences as a function of the number of training CDR-H3 sequences in within-antigen transfer learning. We plotted for each training dataset size the total number of epitopes recognized by generated CDR-H3 sequences for without transfer learning (-T_epitope), with transfer learning (+T_epitope), the total number of epitopes recognized by native CDR-H3 sequences (native_epitope), the overlap of epitopes recognized by both native and generated CDR-H3 sequences for without transfer learning (-T_overlap), and lastly the overlap of epitopes recognized by native and generated CDR-H3 sequences for with transfer learning (+T_overlap). We found that generated CDR-H3 sequences recognized fewer or equal numbers of epitopes compared to the native CDR-H3 sequences. The reverse is true for the across antigens transfer case (Fig. S13).

**Supplementary Figure S13.**
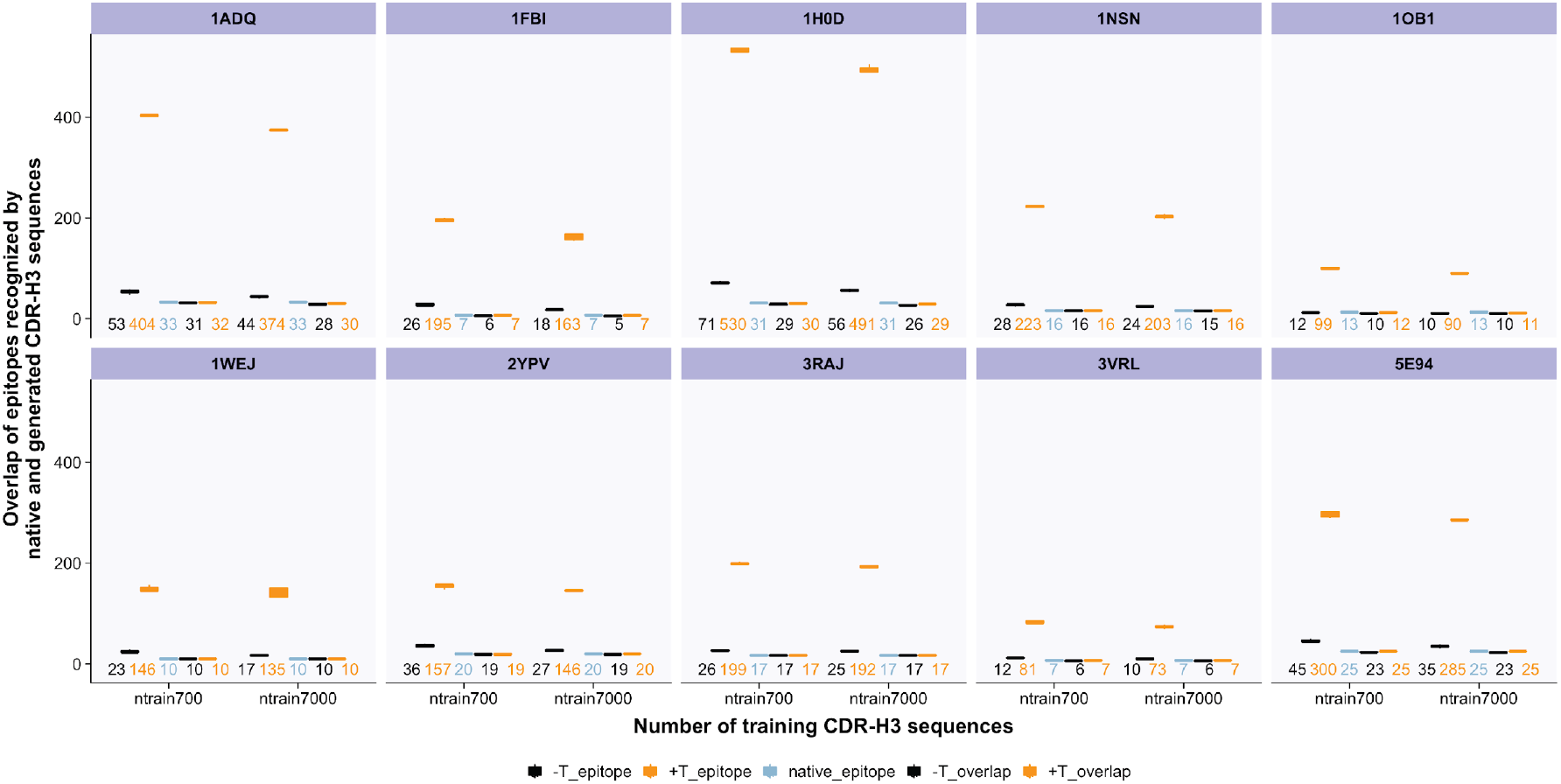
(relates to Figure 4B bottom panel and Fig. S10). Overlap of epitopes recognized by native and generated CDR-H3 sequences as a function of the number of CDR-H3 training sequences in across-antigens transfer learning. To examine the concordance among recovered epitopes in the across-antigens transfer learning setting, we plotted, for each dataset size (ntrain700-ntrain7000), the total number of epitopes recognized by generated CDR-H3 sequences without transfer learning (-T_epitope), with transfer learning (+T_epitope), the total number of native epitopes (native_epitope), and the overlap of epitopes recognized by native and generated epitopes for without transfer learning (-T_overlap) and with transfer learning (+T_overlap). We found that the generated CDR-H3 sequences recognized more epitopes than the native ones.

